# Sifting Through the Noise: A Computational Pipeline for Accurate Prioritization of Protein-Protein Binding Candidates in High-Throughput Protein Libraries

**DOI:** 10.1101/2024.01.20.576374

**Authors:** Arup Mondal, Bhumika Singh, Roland H. Felkner, Anna De Falco, GVT Swapna, Gaetano T. Montelione, Monica J. Roth, Alberto Perez

**Affiliations:** Department of Chemistry and Quantum Theory Project, University of Florida, Leigh Hall 240, Gainesville, FL; Department of Pharmacology, Rutgers-Robert Wood Johnson Medical School, 675 Hoes Lane Rm 636, Piscataway, NJ 08854; Department of Chemistry and Chemical Biology, Center for Biotechnology and Interdisciplinary Sciences, Rensselaer Polytechnic Institute, Troy, New York 12180, United States

**Author notes:** Corresponding authors: Alberto Perez.

**Keywords:** AlphaFold, protein-peptide interactions, protein-protein interactions, BET, BICRA

## Abstract

Identifying the interactome for a protein of interest is challenging due to the large number of possible binders. High-throughput experimental approaches narrow down possible binding partners, but often include false positives. Furthermore, they provide no information about what the binding region is (e.g. the binding epitope). We introduce a novel computational pipeline based on an AlphaFold2 (AF) Competition Assay (AF-CBA) to identify proteins that bind a target of interest from a pull-down experiment, along with the binding epitope. Our focus is on proteins that bind the Extraterminal (ET) domain of Bromo and Extraterminal domain (BET) proteins, but we also introduce nine additional systems to show transferability to other peptide-protein systems. We describe a series of limitations to the methodology based on intrinsic deficiencies to AF and AF-CBA, to help users identify scenarios where the approach will be most useful. Given the speed and accuracy of the methodology, we expect it to be generally applicable to facilitate target selection for experimental verification starting from high-throughput protein libraries.

**Table of Contents:** 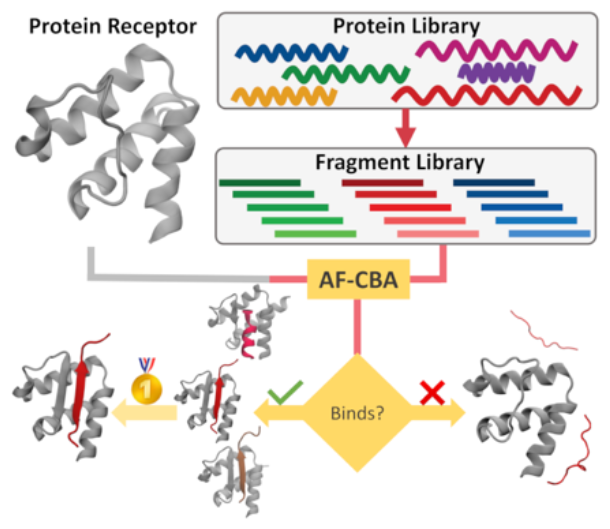

## Introduction

Understanding protein-protein interactions is key to understanding protein functionality, their role in disease, and potential drugability^[1]^. Given the size of the proteome, high throughput methods are essential to narrow the possible network of protein-protein interactions (PPIs) for a target protein of interest^[2]^. However, such high-throughput methods (e.g. pull-down experiments or yeast two-hybrid) produce many false positives, requiring further validation, which is often expensive and labor intensive^[3,4]^. These methods also encounter challenges distinguishing direct PPIs from indirect interactions (see Figure 1A), in which proteins A and B both interact with protein C, but not directly with each other --resulting in a PPI that is not direct^[5]^. Furthermore, these methods do not identify the location in the protein sequence involved in the interaction (see Figure 1B)^[6]^. To understand protein selectivity, characterize binding affinities, or determine the atomic structures of the complexes we would first need to identify directly interacting proteins and their binding epitopes.

**Figure 1.**
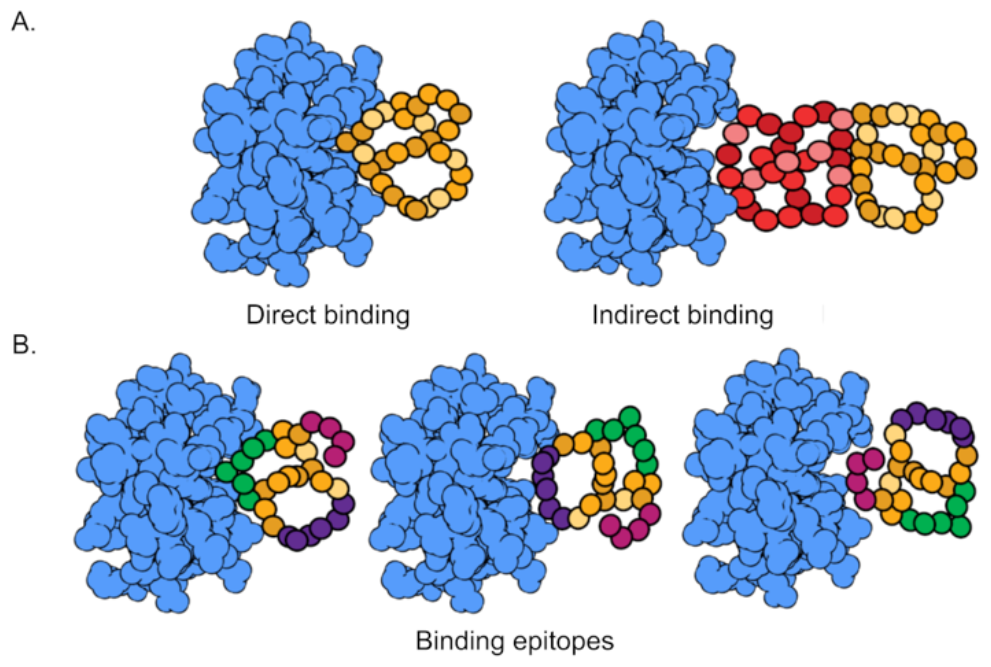
Limitation in high-throughput experiments. (A) A prey protein (orange) can interact directly or indirectly with the bait protein (blue). (B) For proteins in direct interaction, multiple binding modes involving different epitopes (shown here as green, purple, or magenta) are possible, but the experiment will not identify which one is correct.

Computational tools can often help to reduce the number of possible binding partners from a list of candidate proteins. For example, tools such as protein-protein docking, bioinformatic searches using specific binding motifs, or protein-protein interface detection tools have been developed to help identify binding epitopes and, in some cases, the structures of the complex^[7–10]^. Despite progress, there are still many caveats that prevent systematic and accurate identification of binding epitopes^[11,12]^.

In this work, we leverage the recent successes of AlphaFold2 (AF)^[13]^ and an AlphaFold Competitive Binding assay (AF-CBA)^[14]^ to design a novel computational pipeline that identifies the top candidates involved in PPIs, and the location(s) of the protein-protein interaction region(s).

Our efforts are focused on identifying plausible PPIs involving proteins that bind the Extraterminal (ET) domain of Bromo and Extraterminal domain (BET) proteins. BET proteins regulate genes associated with immune response, and each BET protein exhibits preferential expression in distinct tissues^[15,16]^. Their misregulation is involved in multiple diseases and can also be targeted by viruses and bacteria^[16]^. Many of these interactions take place through the ET domain, which binds short continuous peptide epitopes that fold upon binding^[17]^. Notably, the ET domain can undergo substantial conformational changes during binding, allowing for diverse binding modes^[18]^. Structurally, the Protein Data Bank (PDB)^[19,20]^ contains information on six distinct peptides binding to the ET domain^[17]^. Additionally, a recent pull-down experiment on BET proteins has identified a substantial pool of 752 potential proteins predicted to bind to the ET domain^[21]^.

In these pull-down experiments, ET acts as bait and is tagged with an affinity tag for easy detection and purification. The tagged bait is then immobilized in an affinity resin and as the prey proteins (possible binding partners) are passed through the affinity resin some will form complexes that are later eluted using a specific eluting buffer tailored to the affinity resin in use. Analyzing the complexes (e.g., through gel electrophoresis with a subsequence western-blotting techniques or mass spectrometry techniques) yields the network of potential PPIs^[21]^. Conducting structure determination experiments on all possible proteins and candidate polypeptide regions detected from pull-down experiments is labor-intensive and expensive. Given the potential for false-positives and the lack of knowledge of the binding epitope, the likelihood of success is diminished in most cases. Traditionally, researchers focus on characterizing proteins with the highest number of components identified in mass spectrometry^[22]^, leaving other potential binders unstudied. The goal of our computational approach is to prioritize the most promising candidates for direct interaction.

We first show proof of concept of the methodology by recovering the known peptide epitopes from a series of protein-peptide complexes. We then show transferability of the method to nine unrelated protein-protein systems recovering the known binding epitope (see SI methods, and Table 1). Finally, we present a list of 20 proteins from the pull-down experiment that our method predicts bind ET with high affinity, along with the predicted binding epitope. From this list, we experimentally validate one of the predictions that exhibits a previously unknown binding motif.

**Table 1.**
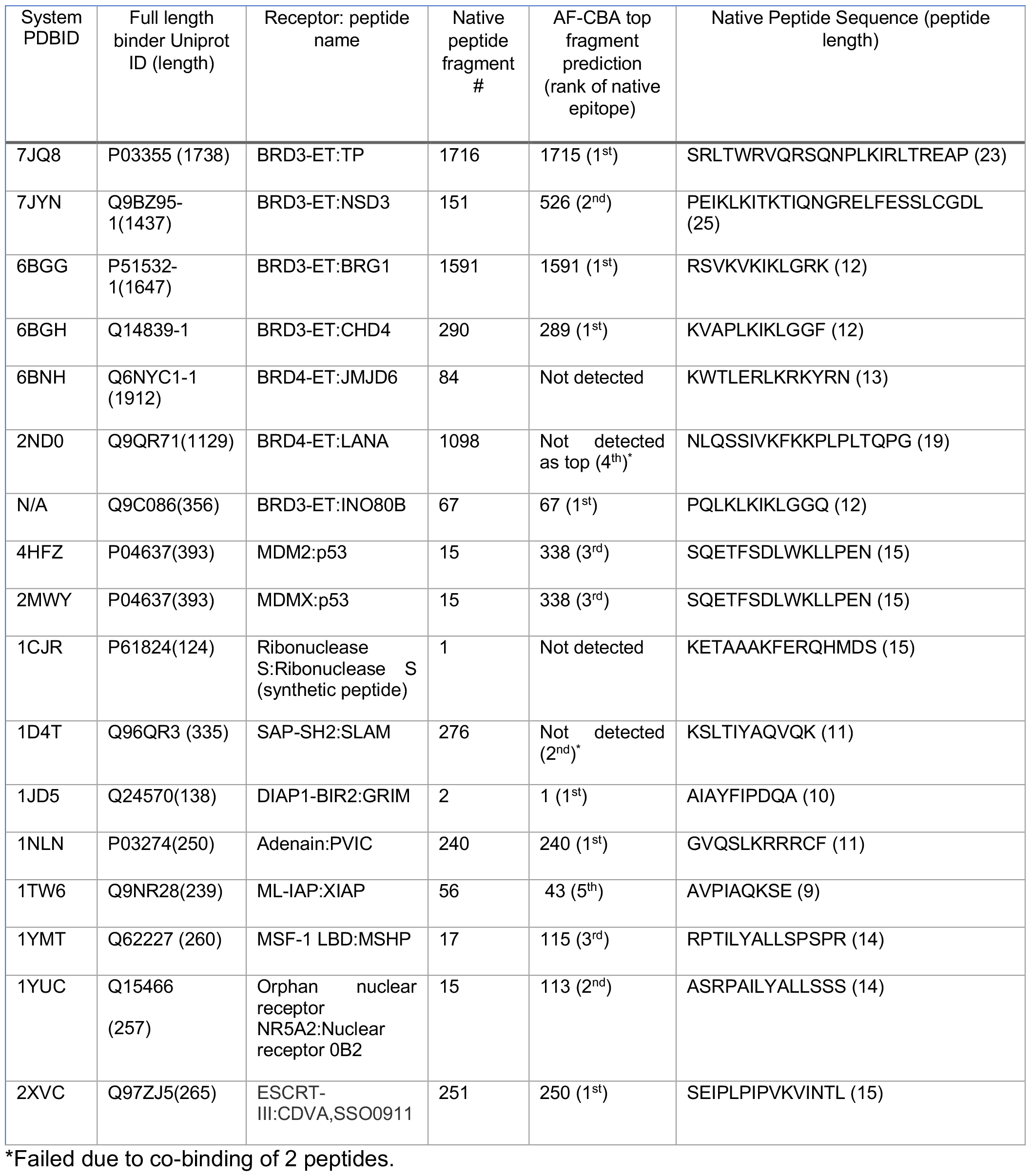
Details of the known systems that we studied in this work.

## Results and Discussion GENERAL APPROACH

Leveraging AF and AF-CBA, we have designed a pipeline (see Figure 2 and SI methods) that starting from a protein sequence goes through three filtering stages to determine if a protein is likely to bind a target and identify the binding epitope. In short, we fragment each protein of interest into a library of overlapping peptide epitopes. In the first filter we run an AF prediction for each peptide in the library to identify if the peptide binds. In the second filter we select five random peptides from the pool of binders and compete each peptide in the pool against each of the five peptides – we select those peptides that outcompete the random peptides most often for the next stage. In the third filtering stage we do an all-by-all competition of all remaining peptides, which leads to identifying the strongest binder in the set. The SI methods contain information about how we select which peptides advance after each filtering step.

**Figure 2.**
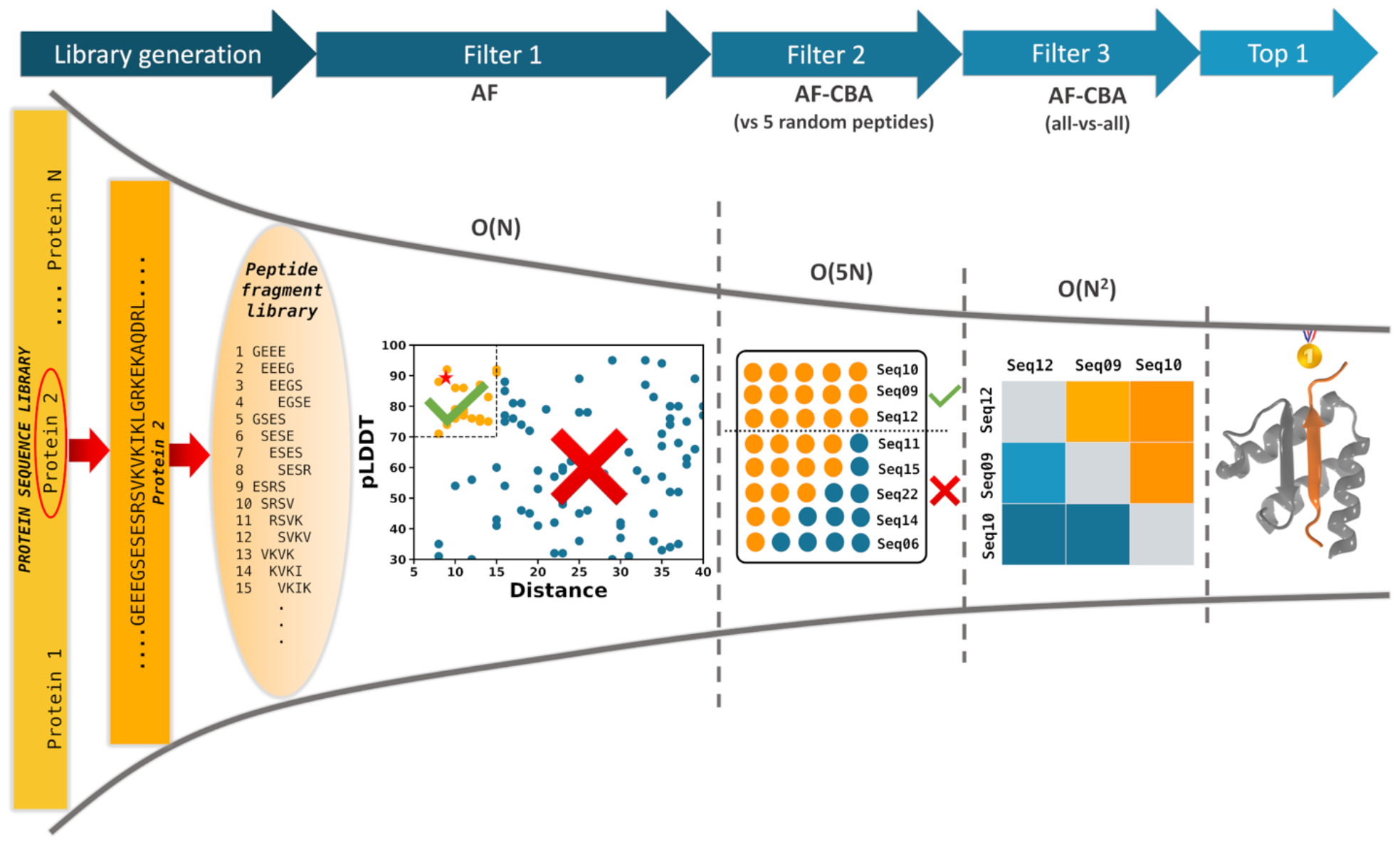
AF-CBA Pipeline. Starting from a library of proteins (left), we create a peptide library for each protein, and run it through a three-filter stage pipeline (from left to right). The first filter is an AF binding prediction for each fragment to the receptor. Top hits are selected based on a combination of pLDDT score and distance of the peptide to the binding site. We select five random peptides from the resulting binders and compete other binders to it using AF-CBA – we select those peptides that most often outcompete the random peptides (filter 2). In the final stage, we do an all-by-all AF-CBA for remaining peptides and select the strongest binder as the one that most often outcompetes all other peptides (filter 3). The computational scaling cost of each stage is provided.

With enough computational resources one could just run the all-by-all comparison, but the cost increases O(N^2^) where *N* is the number of peptides to compete. The initial AF filter only scales as O(N), and the second filter as O(5N). Since N is reduced at every filtering stage this significantly reduces the overall computational cost (see Figure S1).

## PROOF OF CONCEPT 1: RECOVERING KNOWN PEPTIDE BINDERS TO BET PROTEINS

The ET domain is a three-helix bundle with a flanking loop that can adopt multiple conformations to bind multiple host regulatory proteins (as well as viral and bacterial). The flanking loop contains a pattern of alternate negatively charged and hydrophobic residues that constitutes part of the binding domain, together with a hydrophobic pocket formed by the helices^[17,23,24]^ (see Figure S2). Most notably, some peptides interact by forming strand pairing interactions with the loop region of ET and a complementary pattern of positively charged and hydrophobic residues^[17,23]^ (Figure S2). Some peptides bind as a beta-hairpin rather than single strands, completely occupying the hydrophobic pocket in ET^[23]^. One peptide, JMJD6, has been observed to bind using a different structural motif containing an α-helix and lacking the pattern of alternating hydrophobic/charged residues^[25]^.

We took the sequence of seven proteins, including INO80B, a protein with an unknown complex structure (with BRD3) but known binding epitope^[21]^). These proteins are known to bind the ET domain of BET proteins (see *Systems of study* in SI Methods, Figure 3, and Table 1) and created a fragment library for each based on the known epitope length (see *Sequence fragmentation* in SI methods). This resulted in peptide libraries of different size according to the length of the protein and length of the peptide epitope (between 345 and 1901 peptide fragments, see Table 1). We then ran our three-stage filtering pipeline to identify the predicted top binding epitope for each of the seven proteins. Figure 2 exemplifies the filtering process through the pipeline, starting from all possible peptide fragments until the top candidates are identified.

**Figure 3.**
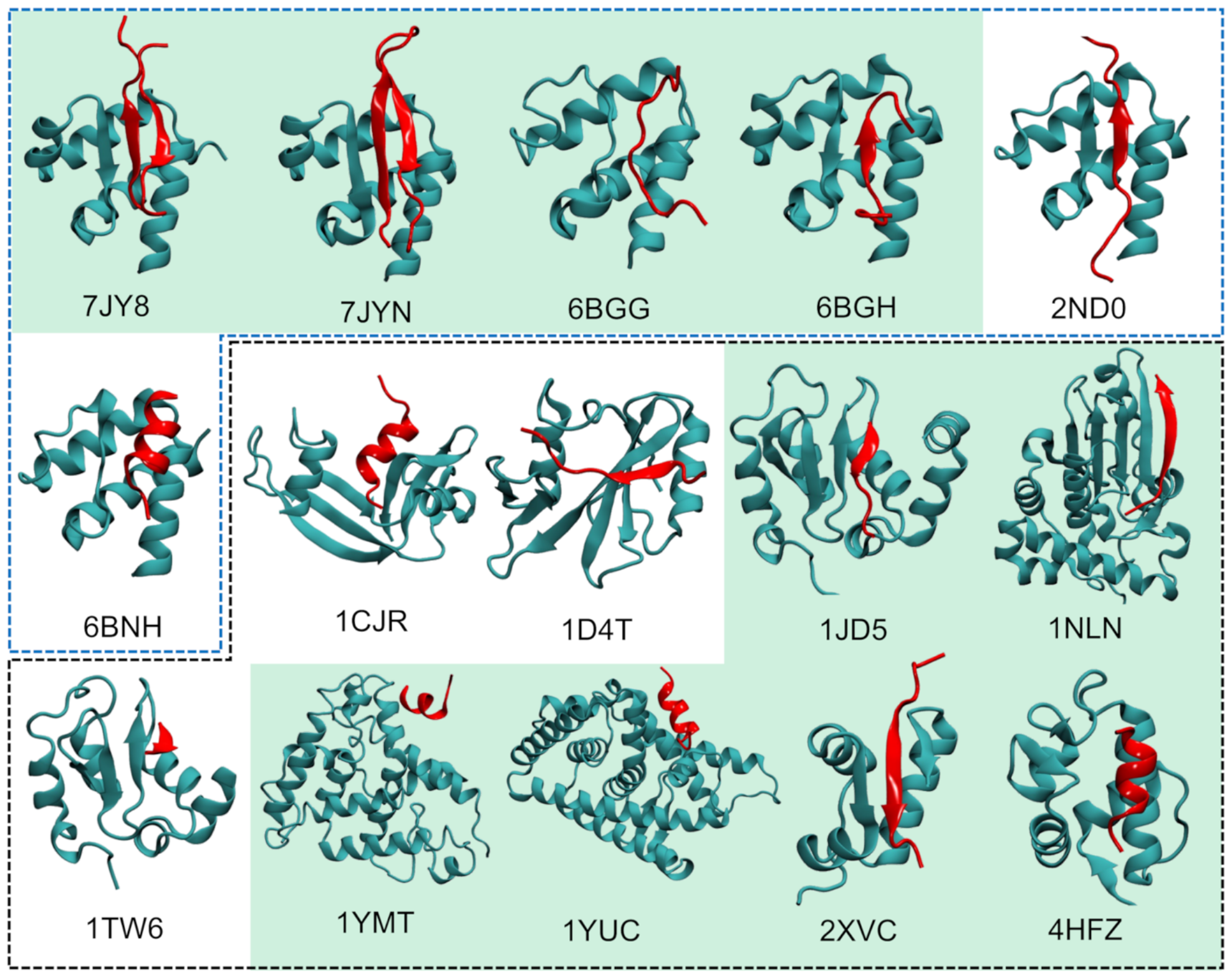
Test cases for AF-CBA pipeline. Experimental structures of protein complexes used in this study with their PDB code. The receptor protein is shown in cyan and peptides in red. The blue dashed line denotes systems involving the ET receptor (Proof of concept 1), whereas those inside the black dashed line show transferability to other systems (Proof of concept 2). Successful cases are highlighted with green background, new insights and limitations are discussed in the text for those with a white background. These representations are rendered with VMD^[26]^.

For four of the systems (TP, CHD4, BRG1, and INO80B), AF-CBA predicts the known binding peptide sequence, plus/minus one amino acid (see Table 1, Figure 4A-C and Figures S3-S6). This potentially means that the core region along those sequences is the one that is indispensable for binding. Indeed, when looking at the NMR structures of TP:ET, BRG1:ET, and CHD4:ET, we find that some of the N-terminal and C-terminal residues of the selected peptide are not interacting with ET and could potentially have been excluded from the NMR study without affecting the binding mode and affinity^[21,23]^. The same pattern is observed for the AF predicted INO80B structure, where fragments 64 to 67 bind through the same set of residues (Figure S5C).

**Figure 4.**
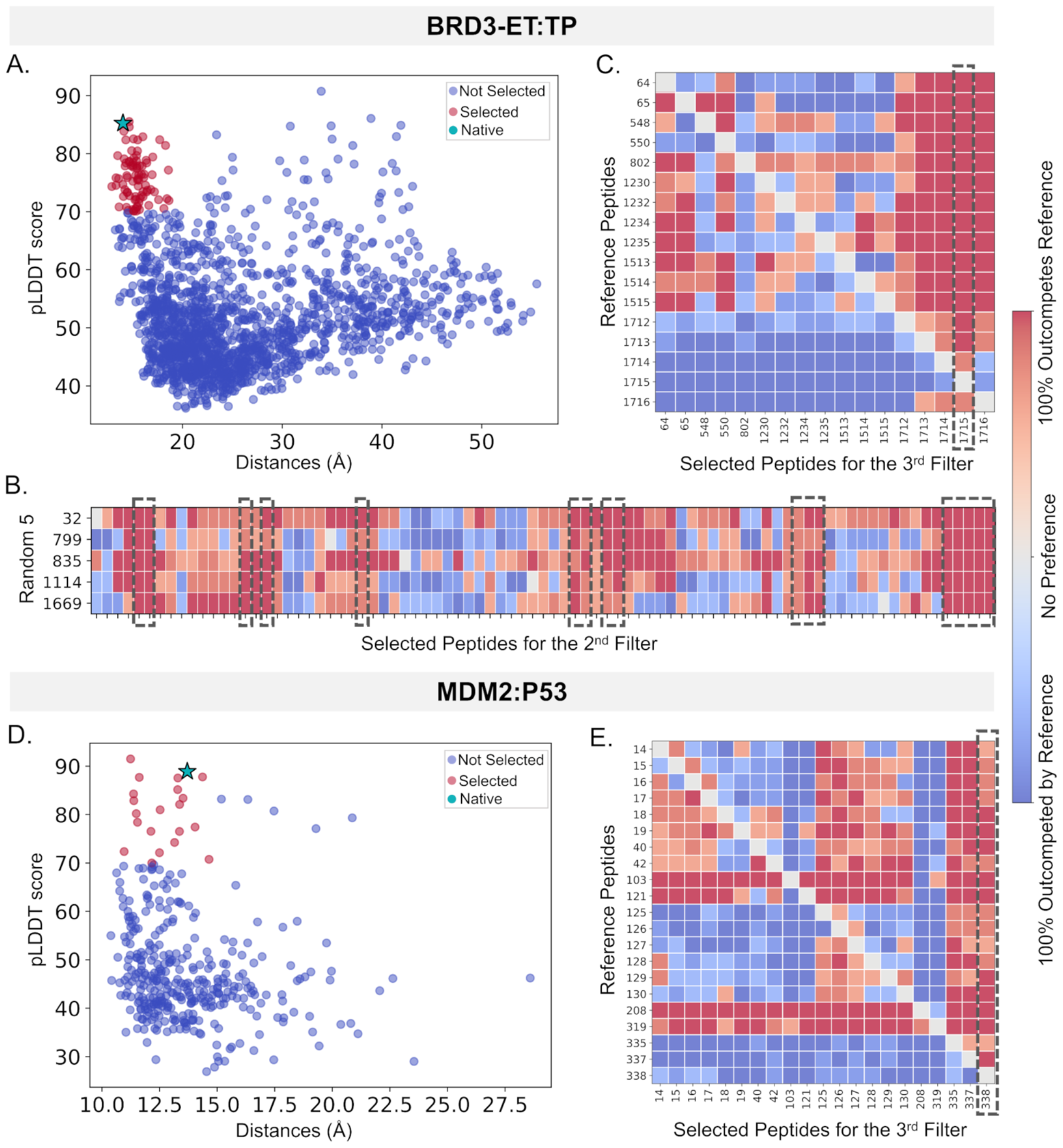
AF-CBA pipeline results for BRD3-ET:TP and MDM2-P53. (A) Filtering AF predictions for all possible epitopes of full length MLV-IN against predicted pLDDT score (y axis) and average distance between all peptide residues to key residues in the receptor ‘s binding site. Each dot is a prediction, with those in red advancing to the next stage, and the experimentally assigned peptide shown as a star. (B) AF-CBA between each peptide candidate in the remaining pool (each column) and each of five random peptides (each row). Dark red columns (see color bar) represent more successful binders and thus advance to the next filtering stage (denoted by a black dashed box). (C) The remaining pool of peptides is evaluated in an all-by-all AF-CBA experiment. Rows denotes peptides used as reference; each column shows the results of each peptide competing against each reference. The column with more red represents the highest ranked peptide binders (denoted by a black dashed box). (D) AF binding study for P53 against MDM2 similar to panel A. (E) Given the small number of peptides after the first filter (<30), we skip the second filtering stage and perform an all-by-all AC-CBA experiment.

When searching for new peptides, we typically don ‘t know what fragment size to use. However, AF might itself help identifying the core residues needed for binding based on overlapping fragments. To test this hypothesis, we created a library of peptides for INO80B considering a length of 25 residues, rather than the 12-residue length known from experiments. Indeed, AF-CBA predicts several peptides that bind with similar affinity, all of them containing at their core the same binding epitope as the shorter, experimental peptide epitope (see Figure S6). With the longer epitopes we observed binding patterns in which INO80B binds as a hairpin (as does NSD3 and TP). Intriguingly, we observe a flipped orientation of the peptide in this case with respect to the shorter (12 residues) fragment ‘s orientation (see Figure S6).

For the NSD3 system, AF-CBA predicts the native sequence as the second-best epitope (see Table 1 and Figure S7). Notably, the best scoring peptide fragment (526), which is also predicted to bind by forming a three-stranded beta sheet, does not have a canonical alternate charge-hydrophobic residues, and binds in an orientation like that of the TP-ET complex. Interestingly, previous experiments have shown several possible NSD3 epitopes that bind ET along the protein sequence, with a range of binding affinities, highlighting the difficulty in ascertaining what the binding epitope should be or even how long the peptide should be^[21,27]^. For instance, while the original NSD3 binding region was identified as a shorter fragment which forms a 2-stranded beta-sheet interaction with ET^[23]^, we found that a longer peptide can form a more stable 3-stranded beta-sheet interaction. However, even this longer this peptide (resides 150 to 184, or Fragment 150 in our study) binds with relatively low affinity K_d_∼250 µM)^[17]^ . Unfortunately, the poor solubility of a synthetic peptide corresponding to this newly predicted ET-binding region of NSD3 (Fragment 526) has so far prevented experimental validation of its binding affinity.

Finally, for the low binding affinity LANA(K_d_∼635µM^[24]^)and JMJD6 (K_d_∼160 µM^[25]^), our pipeline does not select them as probable candidates for experimental validation. The LANA peptide, with a small interaction region and disordered tails, is filtered out of the first stage, as the disordered tails reduce the overall pLDDT below the threshold (Figure S8A). Even after relaxing the filtering criterion (average pLDDT > 55%), AF-CBA predictions exhibit co-binding of the competing peptides to the active site at the 3^rd^ filtering stage (Figure S8B-E), rendering the method unsuitable for this system as it cannot differentiate independent binding of the peptides. For JMJD6, AF fails to predict the native structure of the complex with the ET domain (Figure S9A-B). Due to a very low pLDDT, the native epitope is lost after the first round in our pipeline which is an indication that the assay is not meaningful for this system^[14]^. Generally speaking, for the ET system AF-CBA works better for peptide-protein interactions that are moderately or very tight (K_d_ < ∼ 100 μM) than for more weakly interacting peptides that have been reported previously.

We further delved into JMJD6 and LANA, thinking about our generalizable approach for predicting epitopes in new systems and the known limitations in selecting the epitope for experimental verification. In particular, the JMJD6 epitope was experimentally characterized based on a bioinformatic search for an alternating positive/hydrophobic pattern of interaction. Interestingly, the original paper mentions that a conformational change in JMJD6 is needed for that epitope to be accessible for binding^[25]^. We know, for example from NSD3, that a bioinformatics search can return multiple binding epitopes, all of which bind ET, with different affinities – and not all are relevant to protein-protein interactions. Furthermore, our work showed using a longer NSD3 binding epitope increased binding affinity^[23]^. Thus, we set to try our predictive pipeline, where we select all possible epitopes 25 residues in length. We notice that for JMJD6, the best prediction binds as a hairpin, to form a three-stranded beta sheet, with an orientation similar to that of the ET:TP complex, while the second-best prediction binds as a single strand (Figure S9C-E). Both of these binding modes are more compatible with all other known systems binding ET. However, considering the AF predicted the full length JMJD6 structure, this best predicted epitope may not be accessible for interacting with ET domain (Figure S9E).

For the LANA system, AF-CBA shows co-binding of two strands is favored over a single strand binding (see Figure S8). Such co-binding is reminiscent of when peptides bind as hairpins. Indeed, using 25 residue fragments for LANA we observe binding modes compatible with the expected antiparallel hairpin (Figure S10). In particular the first and second ranked predictions bind as a hairpin with orientation similar that observed in ET:NSD3 complex^[23]^, while the third ranked prediction (containing the previously experimentally determined epitope of LANA) binds in a flipped orientation like that of ET:TP complex. Moreover, an AF prediction of the full-length LANA protein sequence indicates that the best AF-CBA prediction overlaps with a disordered region and thus more likely to be accessible for binding (Figure S10C). Interestingly, for the 25 residue LANA fragment we see many peptides in the first filter screen that bind with high pLDDT scores but far away from the known binding site. While we did not systematically analyze these, visual inspection showed that in some cases these correspond to an unbound prediction in which the peptide is structured far from the protein (Figure S11). The same is true for TP and NSD3.

Hence, AF-CBA pipeline successfully pinpointed five proteins as binders while indicating that for the two low binding affinity epitopes (JMJD6 and LANA) the approach would not recommend further experimental validation. More intriguingly, using longer epitopes suggests changes in binding preferences in the three systems we tried (INO80B, JMJD6 and LANA), indicating that there might be better binding epitopes (e.g. longer variants or a different region) than the ones that have been previously reported in the literature^[21,24,25]^. For LANA and INO80B, the high-ranking predicted peptide epitopes contain the previously detected region, while for JMJD6, AF-CBA predicts an alternate region, with a non-canonical bioinformatic motif which would explain why it was not previously considered as a possible peptide epitope. Among the five binding proteins, AF-CBA accurately identified the previously identified and experimentally validated binding epitopes as the top prediction in four instances and as the second-best prediction in the fifth case. Notably, AF-CBA also suggests a potential novel binding epitope for NSD3, warranting experimental validation. ndeed, the previously identified binding epitope of NSD3, while clearly interacting with ET, has a very weak binding affinity^[23]^. Additionally, these results demonstrate efficacy in capturing the diverse binding modes and protein conformational changes found in ET complexes, tasks that would be challenging for traditional protein-peptide docking software^[28]^. Thus, while traditional approaches have relied on bioinformatic sequence searches to identify binding epitopes, the AF-CBA approach relies on insights learnt from both sequence and structural data, allowing us to better prioritize epitopes for experimental characterization.

## PROOF OF CONCEPT 2: TRANFERABILITY TO OTHER PEPTIDE-PROTEIN SYSTEMS

AF-CBA pipeline has successfully identified epitopes binding the ET domain. Here, we aim to assess its broader applicability to other systems compatible with AF and AF-CBA. We examine nine additional systems^[29–38]^ (see Figure 3, 4D-E and Figures S12-S20, and Table 1) chosen for their diverse topologies and binding modes. The methodology, detailed in the SI Methods, consistently identifies the native epitope three times as the top hit (+/-1 residue) and within the top three hits in three instances. Here, we consider each region along with its multiple overlapping fragments (generally shifted by one or two residues) as a single hit, thus the second hit would be a region that does not contain these overlapping fragments. Finally, our investigation reveals important limitations in three other systems, as discussed in the limitations section.

For systems 1NLN, 2XVC, and 1JD5, the predictions were highly successful (Figure S12-S14). In all three cases the peptide binds through a beta strand that pairs with another strand in the receptor. Conversely, for MDM2-p53 (Figure 4D-E) and 1YMT, AF-CBA the peptide binds as a helix. For these two systems, our predictions identified the native binding fragment as the 3^rd^ best prediction and, for 1YUC (another helical binder), as the 2^nd^ best (Figure S15-S17). AF has a known bias towards stabilizing helices, occasionally leading to predictions folding peptides into helices that might not be inherently stable. Upon further inspection, for the p53-MDM2 interaction the top prediction contains a similar pattern of hydrophobic residues inserting into the cavity of MDM2 as the native epitope. Upon looking at the full length p53 structure, the second fragment corresponds to a region predicted to be inaccessible (Figure S15A-B) and thus unlikely to bind. We tested the binding of p53 with MDMX receptor which resulted in similar prediction as MDM2 (Figure S15C). We noticed a similar pattern when we investigate the position of the top epitopes with respect to the full-length proteins for 1YTM and 1YUC, which are mostly accessible and presenting a similar pattern of hydrophobic interactions as the solved structure (Figure S16-S17) implying other possible binding epitopes.

The three remaining systems (1CJR, 1TW6, and 1D4T) did not successfully identify the proteins as binders or their binding epitope (Figure S18-S20). The current methodology works within the boundaries of its constituent AF and AF-CBA intrinsic limitations. Identifying systems where these limitations are apparent allows us to exclude the system from further analysis, as the predictions will be flawed. Some limitations lead to overlooking potential binders which leaves the user with no additional knowledge. For example, AF might not predict a discontinuous peptide binding to the receptor unless a small continues region of the peptide contributes strongly to the free energy. Other limitations might confound the ranking when not taking in consideration (e.g., structures where both peptides co-bind during AF-CBA should not be considered for ranking). We discuss these limitations in detail later in the Limitations section. For 1CJR, AF fails to predict the complex accurately, as reflected by a low pLDDT score, and appropriately filters it out at the first stage as same JMJD6 (Figure S18). For 1TW6, the small interaction region between the protein and peptide (4 residues) confounds AF and leading to less accurate prediction of the top binders (Figure S19). For 1D4T, as in the case of LANA, we observe co-binding, rendering AF-CBA ineffective for this system (Figure S20).

Collectively, these two proofs of concept provide insights into the capabilities and limitations of the method. The AF-CBA pipeline assesses individual peptides without contextual knowledge of their position within the protein and considering continuous epitopes. Some epitopes identified by AF-CBA may never be accessible for binding or may necessitate substantial conformational changes in the protein, making them inherently less likely to be involved in PPIs. This challenge mirrors the complexity observed in the promiscuous binding of transcription factors to regulate gene expression. With millions of high affinity binding sites along the genome, many remain inaccessible, resulting in less than 1% of potential binding sites being occupied^[39]^. Additionally, as with peptide epitopes, the length of flanking DNA affects binding affinity^[39]^. In the context of this work, while the top ranked peptides may indeed bind and aid in identifying receptor selectivity, they may not necessarily be biologically relevant as epitopes mediating PPIs. Additional considerations, such as accessibility, play a crucial role in pinpointing the correct epitopes.

## PREDICTIONS: AF-CBA IDENTIFIES THE STRONGEST BINDERS FROM PULLDOWN EXPERIMENTS

The pull-down experiment for ET contains a library of 752 proteins ranging in size from 69 to 4128 residues^[21]^. We fragmented each protein into a peptide library and processed each library as described in the methods section by competing against TP and NSD3. The pipeline identifies 22 proteins that bind ET with high confidence (outcompeted NSD3 at least in 80% of the AF-CBA models, see Table S1). Amongst all systems, we only found one instance (RPA1) that systematically outcompetes TP. The TP peptide epitope is the C-terminal segment of the Murine Leukemia Virus (MLV) integrase protein whereas NSD3 is a host regulatory protein^[23]^. The particularly high binding affinity of the TP epitope for ET could have evolved to outcompete all host regulatory proteins facilitating efficient, targeted integration into the host genome.

For these high confidence hits, we repeated the fragmentation approach using a one residue sliding window (as we did for the proof of concept) to find the best interacting regions. Interestingly, NSD3 (WHSC1L1) and INO80B also show up in the hit list, however we left these hits out as we had already tested them as a known system. Table S1 describes the details about these systems along with the sequence of the top one to three predicted regions. Additionally, the AF predicted complex structures of the best predicted regions of these hits with BRD3-ET are shown in Figure S21. Many of these predicted epitopes do not contain the canonical pattern of alternating charged and hydrophobic residues. We further investigated the accessibility of the predicted regions. Since the majority of these full-length proteins were not solved experimentally, we used AF predicted structures from the AF database (https://alphafold.ebi.ac.uk)^[40]^. Figure S22-S23 show all the full-length predicted protein structures, with the predicted binding regions highlighted using different colors. Based on these structures, some of the predicted epitopes may not be accessible for interacting with ET. While our method significantly narrows down the number of candidates from the original 752 proteins, other considerations such as biological implications from GO terms^[41]^ might further help to prioritize binders for experimental verification.

Amongst the list of possible binders, GLTSCR1, also referred to as BICRA (BRD4 Interacting Chromatin Remodeling Complex Associated Protein) is a 1560-residue protein that as the name implies is already known to bind BET proteins (Uniprot ID: Q9NZM4)^[27,42,43]^. Its detection as a binding partner provides a validation of the AF-CBP method. However, it is unknown which region of BICRA is the binding epitope or how it binds to ET. We looked at the best three predictions from our pipeline and mapped them to an AF prediction of the BICRA protein in isolation to verify that all three epitopes correspond to regions predicted to be flexible and accessible for binding (Figure 5A-D). The two first epitopes bind with an NSD3 orientation, while the third binds with a TP-like orientation (flipped hairpin).

**Figure 5.**
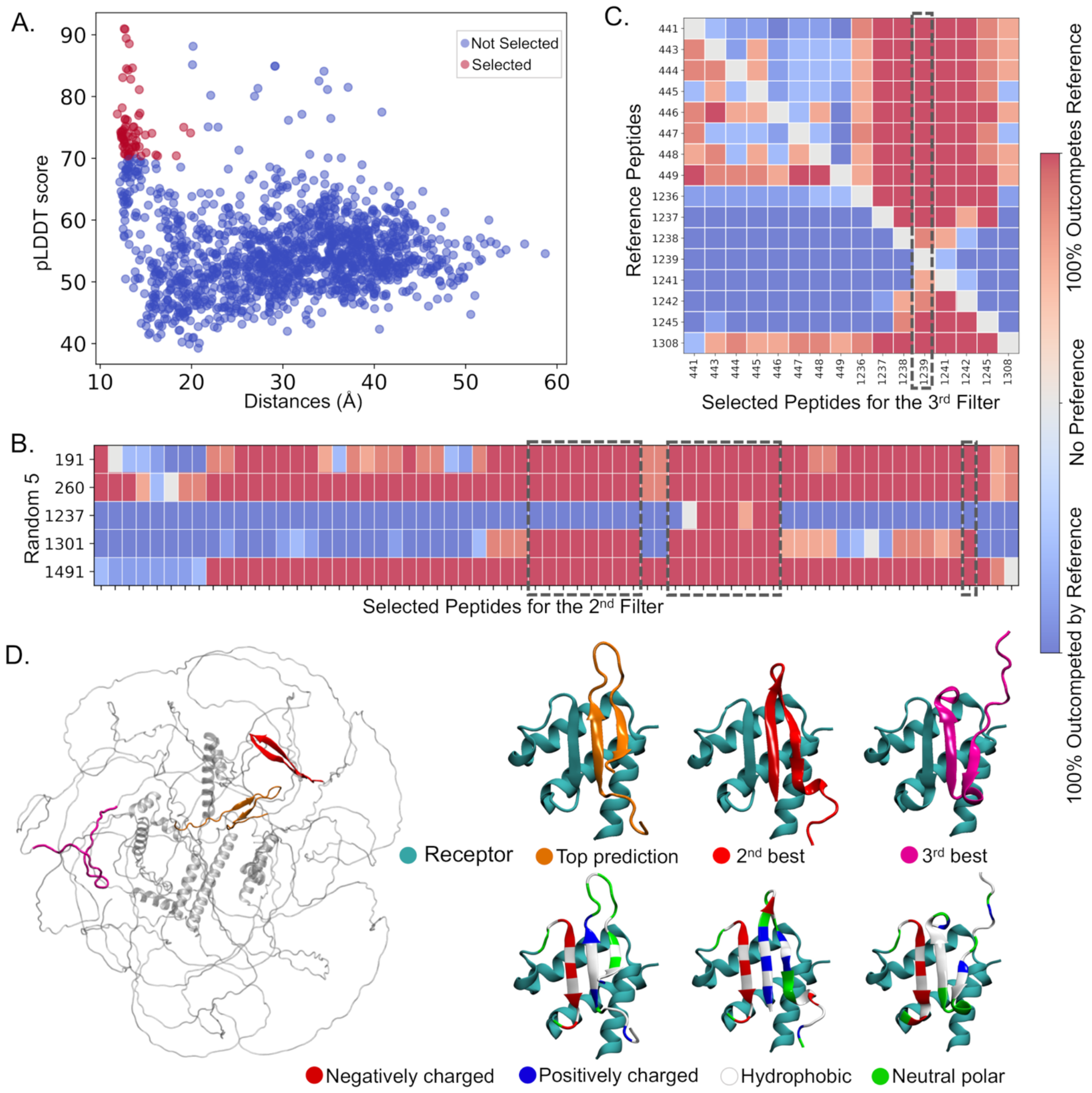
AF-CBA prediction and experimental validation for BRD3-ET:BICRA. (A-C) Three stage AF-CBA pipeline result similarly represented as in Figure 4. (D) AF predicted full length BICRA with three best binder regions highlighted (left) and complex structures of top three predicted epitopes with BRD3 (right) where only the 2^nd^ best prediction shows the canonical alternate opposite charge zipper.

What is unusual about the top (fragment 1239) AF-CBA predictions is the lack of an alternating pattern of hydrophobic and charged residues typical of ET binding peptides (Figure 5D). Thus, this epitope would not have been picked up by traditional bioinformatic approaches or human intuition. On the other hand, the second prediction peptide region (fragment 1308) does present an “LKLKIK” pattern of hydrophobic and charged residues which human intuition and a bioinformatics search would have selected from the protein sequence^[23,25,27]^. Indeed, a neighboring peptide (fragment 1298) was suggested in earlier literature based on a bioinformatic search^[21]^. We had previously extended this reported peptide to incorporate a motif capable of binding as a hairpin (fragment 1304).

## EXPERIMENTAL VALIDATION OF PREDICTED ET-BINDING POLYPEPTIDES

We developed an assay in which each peptide of interest was synthesized coupled with an Ahx linker to a biotin molecule at the N-terminal end (see SI methods and Figure 6A). The His_6_-tagged BRD3 ET domain was immobilized to nickel-coated plates through the His_6_ moiety. Biotin-conjugated peptides of interest were then titrated for binding with molar ratios ranging from 0.01 to 160-fold excess with respect to the bound ET (9 pmol). Bound peptides were quantified through conjugation of strepavidin poly-HRP to the peptide-linked biotin and measuring HRP with visual light (λ_350-700_ nm). Figure 6B shows these results as binding saturation curves to TP and to proposed peptide epitopes of BICRA. Surprisingly, the top ranked AF-CBA prediction for BICRA peptide binding epitope is indeed a stronger binder than peptides containing the more typical alternating charge binding pattern (see Figure 6B). This is an unexpected success of the AI pipeline which has learned beyond the specific patterns of interaction expected of ET.

**Figure 6.**
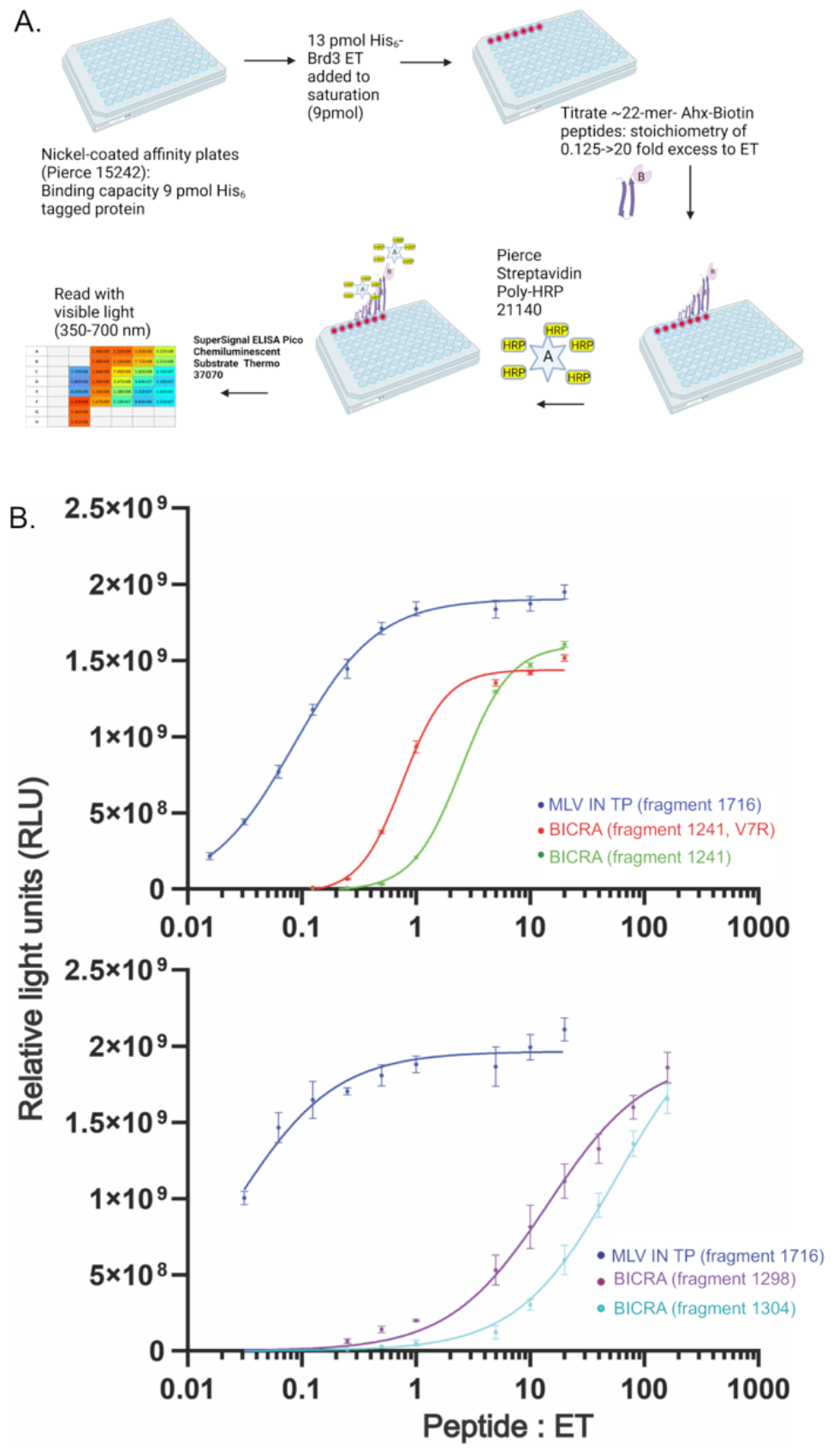
Experimental validation of BRD3-ET:BICRA. (A) Schematic representation of the experimental procedure. (B) Binding curve for the best predicted epitope of BICRA, V1247R mutated epitope, and native TP against BRD3 (top) along with 2 epitopes near second best predicted region and TP (bottom).

While the BICRA polypeptide epitope was found to be a strong binder, it is not as strong as the Tail Peptide from MLV-IN, the strongest known binder to ET^[23]^. We introduce a V-> R mutation in the top binding sequence ( “KLVIR “) to recover the recognizable pattern of alternating charged/hydrophobic residues. Indeed, experiments show that the new sequence is an even stronger binder – but not yet at the level of TP (Figure 6B).

## LIMITATIONS

We have found 3 different scenarios where the current approach does not allow the user to prioritize binders. In the first scenario, AF was not able to predict the structure of the complex correctly. The short-length JMJD6 epitope and 1CJR are two instances for this (Figure S9 and S18). Fortunately, we could readily see this from the low pLDDT score, hence, the method is aware of systems for which it will not work.

A second limitation is when the peptide makes too few contacts with the receptor or the binding epitope has low binding affinity. In 1TW6, the x-ray structure of the epitope contains only four residues (Figure 3 and Figure S19). Such low number of contacts is most likely confounding AF ‘s ability to differentiate with its learnt potential. Similarly, AF-CBA finds it hard to predict the native epitope for 1YMT (3rd best prediction is native) where a small alpha helix interacts with the receptor (Figure S16).

The third limitation arises when AF-CBA finds co-binding of both peptides as in 1D4T (Figure S20). In this case, AF correctly predicts the structure of one peptide binding the receptor, but during the AF-CBA both peptides co-bind resulting in a “flipped” binding mode (Figure S20). This was also observed in the case of LANA. Systems with this issue cannot be ranked by this approach.

Our current approach considers contiguous peptide epitope binding fragments. However, some protein-protein interactions, are mediated through different regions along the sequence that are close in space and interact together in the binding site^[44]^. The current approach as described would not be useful in such cases – it might at best identify the independent contiguous fragments that bind occupying different binding modes. For instance, for LANA, when two peptides co-bind, they exhibit a binding mode occupying the whole ET binding site as other peptides that bind as hairpins would do. This led us to look for longer LANA fragments and indeed recovered a stronger binder that binds as a hairpin. In other scenarios the different secondary elements might not be close in sequence space but still interact together in the binding site.

## SYNERGIES WITH OTHER APPROACHES

We have highlighted synergies with experimental high-throughput data and the ability to prioritize which epitopes are more likely to warrant follow-up experimental characterization (e.g., binding affinities and/or structures). At the time of writing two other computational approaches have emerged focusing on similar desires to identify binding epitopes. One approach uses a similar fragmentation strategy to identify how peptides bind AF (single protein-peptide binding)^[45]^. As reported in our work, epitopes that are shared amongst overlapping fragments tend to experience the same behaviour (e.g., binding). In their case, they looked for the highest number of overlapping fragments to identify the binding epitope. While this strategy has merit, it can in some cases not be adequate as multiple effects such as epitope length can confound AF. Our AF-CBA assay eliminates this ambiguity by making a head-to-head comparison of the different plausible epitopes. In their work, pre-calculated MSAs, accelerate predictions, which could be combined with our current approach. However, the ability for AF to rank peptides through AF-CBA does not require MSAs for the peptide. A second approach^[46]^, highlights the increased AF sensitivity when predicting PPIs using small protein fragments. In their extensive benchmark they show cases where the whole length protein confounds AF, reducing in lower accuracy predictions or even impeding binding. Their work, validated with experiments, also shows the reduced selectivity for AF over different possible epitopes. Indeed, they show that extending the length of the epitope might result in improved interactions and identification of more complete epitopes than predicted in the absence of a computational technique. However, they warn that extending the size of the peptide is not a general answer as non-interacting tails can sometimes interfere with binding predictions.

## CONCLUSIONS

We have introduced an AF-CBA pipeline that identifies the most likely binding epitope out of a library of peptide fragments. The method works well for peptides that bind the ET domain, where we have shown that it reproduced known binders and have verified one of its predictions experimentally. We have further shown that the method is transferable to other protein-peptide systems.

We expect the rules of thumb for successful AF and AF-CBA predictions will carry for this new assay: 1) binding is mediated primarily by a single continuous polypeptide epitope; 2) high pLDDT scores for structure predictions, 3) peptide adopts well defined secondary structure; 4) medium to strong affinity binders; 5) peptides should not co-bind to the receptor; 6) low sensitivity to minimal mutations. We expect this methodology to be of general interest to understand PPIs, identify binding epitopes, and search/design novel epitopes with higher binding affinity.

## Supporting Information

We describe the experimental and computational pipeline in the SI methods and add supplementary figures to support our results and discussion.

## Acknowledgements

This work was supported by National Institutes of Health grants R35 GM122518 (to M.J.R.), R35 GM141818 (to G.T.M.), and R01GM149646 (A.P.).

## Data Availability

A README file and the scripts used in this work can be downloaded from https://github.com/PDNALab/AF-CBA-pipeline

We introduce a computational pipeline that identifies peptides epitopes that bind a particular receptor from a library of possible protein sequences (e.g., coming from high-throughput experiments). The pipeline uses an AlphaFold-Competitive Binding Assay (AF-CBA) to rank order different plausible epitopes and select the most likely one.

Institute and/or researcher Twitter usernames: @UF, @UFChemistry, @Al Perez, @amondal_chem, @bhumika_chem, @rpi, https://www.linkedin.com/in/gaetano-montelione-0346833

