## Supplementary material for "Sifting Through the Noise: A Computational Pipeline for Accurate Prioritization of Protein-Protein Binding Candidates in High-Throughput Protein Libraries": SI methods and figures

##### Systems of study

For each system we start with the full-length protein sequence, with the aim of identify: 1) if they bind the receptor, and 2) the binding epitope. We have two parts in our study: one where AF-CBA is tested on 17 known systems (Figure 3 and Table 1) and in another part, AF-CBA was used to characterize unknown binder of BRD3:ET from a pull-down experiment (Figure S21-S23 and Table S1).

Systems with known binding epitopes (proof of concept):

*BET protein family:* we tested on the known ET interacting proteins for which we have NMR structures: Tail Peptide (TP) from Murine Leukemia Virus, LANA from Kaposi's sarcoma virus, and host regulatory proteins: NSD3, BRG1, CHD4, and JMJD6. We also included INO80B, where the binding epitope has been reported in the literature, but no structure is yet available. In accordance with the solved experimental structures, we use BRD3-ET as receptor for TP, NSD3, BRG1, CHD4, and INO80B – and use the homologous BRD4-ET for LANA and JMJD6.

*Transferability to other PPIs*: Figure 3 shows the known systems that we tested on. The set was designed to contain a high degree of structural diversity in terms of peptide structure and their binding sites. We selected these systems from PeptiDB[1] and PPDbench[2] protein-peptide databases and used the RCSB reported receptor sequence for these systems. We then obtained the full-length protein sequence using the *scan prosite* server using the peptide fragment as the search motif (https://prosite.expasy.org/scanprosite/)[3].

Potential binders with unknown binding epitope (predictions):

*Potential binders to BET proteins from pull-down experiments:* The 752 proteins from the pull-down experiment represents our protein library. For each protein in the library, we retrieved the protein sequence using their gene name and using the *esearch* command from the *edirect* tool[4]. This resulted in multiple sequence hits for each gene name, and we chose the first sequence that is associated with human. We then used BRD3-ET as the template receptor in accordance with what was used in the pull-down experiment.

**Sequence fragmentation: generating a peptide library from the protein sequence**

To generate a library of peptides, we first need to identify different peptide fragments. We control the library through two parameters: peptide length and sliding window size. A sliding window of one guarantees we generate all possible fragments of a certain size – but increases the number of calculations. We operate in one of two ways for our proof of concept (where we have few protein systems) and for our predictions (where we have a large library of proteins).

For our proof of concept using known binders, we set the peptide length to the known length of the peptide and a sliding window of one. The only exception was NSD3, for which the 34-residue fragment includes a disordered tail which does not contribute to binding – thus in this case we settled for a 25-residue peptide length. For a protein with $L$ residues and a length of 25 residues, this approach will result in L-24 peptide fragments.

For predictions from pull-down experiments, where the number of proteins in the library is large (752 proteins[5]), the previous approach is computationally demanding. We have the advantage of the accumulated knowledge on this system based on known binders and their affinities. Thus, our intent is to discover new possible binders that are relatively strong binders so they can easily be characterized by techniques such as NMR. Instead of using AF to identify peptides that might bind (O(N) cost), we conducted AF-CBA for each peptide, comparing against both the TP peptide (nanomolar binder) and NSD3 (micromolar binder) (resulting in O(2•N) computational cost).

Recognizing that much of the binding affinity can be attributed to a few residues in the peptide, we decided to use a standardized peptide length of 25 residues – which accommodate peptides that bind as a hairpin and those that bind as single strands or helices. Furthermore, we adopted a 13-residues sliding window, so that there was enough coverage of all epitopes. Consequently, a full-length protein of length $L$ yielded (L−24)/13 peptide fragments. For proteins in which at least one peptide ranks higher than NSD3 with high confidence, we repeat the analysis using a 1-residue sliding window (for higher sensitivity) to find the best epitope, following the same approach as we did for known binders. In the repeated stage, all the selected (14 very high confidence hits) proteins were fragmented keeping the peptide length to 25 except for SUPT16H. By looking at the initial prediction of SUPT16H, ACTR1L, and SMARCA5, we realized it binds as a single beta keeping long unstructured terminal (Figure S20). Therefore, at this stage, we used a peptide length of 12 residues to generate fragment library.

**Epitope selection: use a three-stage filtering approach using AF and AF-CBA.**

All AF calculations were carried out with ColabFold[6] with its locally installable *localcolabfold* version (https://github.com/YoshitakaMo/localcolabfold). Each AF and AF-CBA prediction takes about 5 minutes to complete. Each AF calculation is done on a single peptide and receptor, helping to identify potential binders. Each AF-CBA is done on a pair of peptides and the receptor, providing a qualitative ranking between the two peptides – the more times the AF-CBA calculation is run for a pair of peptides, the higher the statistical significance for the ranking. A naïve all-by-all approach to identify the best binder in the peptide library leads to a large computational cost (e.g. $N^2$•2, where *N* is the number of peptides in the library and 2 is the number of times the AF-CBA calculation is done). This computational cost quickly becomes prohibitive when considering many possible proteins (Figure S1). However, we expect most peptides in the library to be weak binders at best, which can be easily discarded, reserving the calculations with higher statistical significance for a subset of strong binders. Thus, we designed a computational pipeline consisting of three filtering stages to lower the overall computational cost (Figure 2, S1).

The first filter uses AF to predict how each peptide binds the receptor. We use multiple sequence alignments (MSAs) for *both* the receptor and the peptide. We then measure the distance between key residues in the receptor's binding site and all peptide residues using MDTraj[7]. To choose the key residues, here we looked at the native structures of the complex and picked 4-5 residues from the receptor's binding site. For an unknown system, however, we can look into few AF predicted complex to get insights about the binding site and key residues. We combined this distance metric and the peptide pLDDT score of the top AF model to make a binary decision between binders and non-binders. We set a 15Å distance cut-off for peptides under 20 residues and of 20Å for longer peptides. The pLDDT cutoff is set to 70 for all cases (Figure 4A, 5A). Typically, too many peptides remain in the library to attempt an all-by-all AF-CBA at the end of this stage, requiring a second filtering stage.

In the second stage we select five random sequences from the *N'* remaining peptides. And we run AF-CBA in duplicate for each peptide against each of the five peptides (N'•5•2 calculations). For each peptide we track how many times they have outcompeted each of the random five peptides. We select peptides that outcompete all five random peptides in at least 80% of the AF-CBA predicted models (Figure 4B, 5B). The dashed rectangular boxes represent the selected peptides for the next filtering round. If at the end of the second filtering stage more than 25 peptides remain in the library, we select five new random peptides and repeat this step.

Since the choice of five peptides is random, we could have scenarios where we select five weak binding peptides, which would not reduce the size of the library. A second possibility is that by sheer chance, one of the five random peptides is already one of the top binders – preventing any other peptide from outcompeting all five peptides. It is easy to identify this pattern and circumvent the issue by shortlisting peptides that outcompete the other four peptides and the top peptide from the random set. For example, in the case of BICRA, one of the random 5 peptides is "Sequence 1237" (involving residues 1237 to 1251 from the whole length protein sequence) which is just one residue shifted compared to the top binder. As a result, we can see in Figure 5B, a very few peptides successfully outcompeted this reference peptide. Therefore, to select peptides for the next filtering round, we just considered the competition against the remaining four random peptides.

Once the second stage is completed, we are ready to identify the strongest binder in the set through a third filtering stage. With 25 or less peptides in the library, we can now run an all-by-all AF-CBA calculations (Figure 4C, 5C). Each calculation yields 5 models, and is repeated 2 times, giving 10 models to determine the rank based on how many times one peptide outcompetes the other. This stage requires N''•N''•2 calculations, where N'' is number of peptides at this stage. Gathering all results, we can set a ranking for the top epitope binders. The dashed rectangular box in Figure 4C and 5C represent the top binder as this peptide outcompeted all others. As an additional step, we look into AF predictions of the whole length protein to identify whether the selected epitopes are in generally accessible regions of the protein (e.g., disordered regions or surface) as shown in Figure 5D.

This layered pipeline greatly reduces the overall computational cost from $O(2 \cdot N^2)$ to $O(N+2 \cdot 5 \cdot N'+2 \cdot N''^2)$. Figure S1 shows a three order of magnitude reduction in computational cost when considering long proteins.

**Experimental validation of BICRA epitope predictions**

Materials: Nickel Coated Plates (white, 96-well, # 15242) and Streptavidin Poly-HRP (StpHRP #21140), were purchased from Pierce. SuperSignal ELISA Pico Chemiluminescent Substrate Kit, (# 37070) was purchased from Thermo Scientific. Bovine Serum Albumin (BSA, heat shock fraction, # A7906-100G) was purchased from Sigma-Aldrich Bovine Serum Albumin. $His_6$-BRD3 ET was purified as previously described[8]. Peptides modified at the C-terminus with Ahx-Biotin were chemically synthesized by Genscript.

BRD3 ET : peptide binding assay.
Peptides-Ahx-biotin were resuspended in $H_2O$ for a stock solution of 2 mg/ml using data supplied by the manufacturer. Briefly, the mg of target peptide was calculated with the formula: (mg gross weight of the synthesized peptide) x (% net peptide content) x (% purity by HPLC) =mg of target peptide. Fresh dilutions of each peptide were made into ET buffer the day of each assay and maintained on ice. Peptide were serially diluted to a molar ratio between 0.015625-160X to ET (9 pmol).

13 pmol of purified $His_6$-BRD3 ET in 25mM $Na_2HPO_4/NaH_2PO_4$ buffer, pH 7.2, 100 mM NaCl, 1mM TCEP (ET buffer; 200 mL) was placed in each well of the Pierce nickel coated 96-well plates (9 pmol His-tagged protein capacity) and incubated at room temperature for > 1h with gentle shaking (80 rpm). Unbound $His_6$-BRD3 ET was removed, and plates washed twice with 200 mL ET buffer. 200 mL of each diluted peptide in ET buffer was added to appropriate well, covered with adhesive polyester film and incubate at room temperature $\geq$1h with gentle shaking (80 rpm). The peptide solution was removed from the wells and wash two times with 200 mL ET buffer. StpHRP was diluted to 1:2500 in ET buffer + 1% BSA, covered with adhesive polyester film and incubated at room temperature for 30' with gentle shaking (80 rpm). The StpHRP solution was discarded from each well and wash twice with 200 mL ET buffer. 200 mL Chemiluminescent substrate to each well and read on a Promega GloMax Discoverer plate reader on Luminescence channel between 1-5' after adding substrate. Experiments were performed in triplicate for each peptide concentration. Control samples omitted the peptide-Ahx-biotin in the reaction.

The experimental procedure is graphically summarized in Figure 6A. The outliers were identified using Grubbs' test (alpha 0.05) for all the number sets. The data was plotted using GraphPad Prism version 10.1.2 for Windows, GraphPad Software, Boston, Massachusetts USA, www.graphpad.com. Figure 6B shows the plotted data for 5 systems in two panels. For the systems that plotted in the top panel of Figure 6B, the best fit was sigmoidal with the four-parameter logistic curve, abbreviated 4PL, whereas for the bottom panel systems, the best fit was hyperbola.

**Table S1**. Details of AF-CBA study on the top hits from the pull-down experiment. Interacting residues in the best predicted epitopes are highlighted with yellow.

| Gene | Uniprot ID (Protein length) | AF-CBA top predicted regions | peptide sequence | Accessibility of the best prediction |
|---|---|---|---|---|
| SUPT16H | Q9Y5B9 (1047) | 646,<br>355,<br>716 | VKQDSLVINLNR<br>EFREGSLVINSK<br>MIIVLHFHLKNA | Yes |
| RUVBL2 | Q9Y230 (463) | 191,<br>143 | DVITIDKATGKISKLGRSFTRARDY<br>VEIQIDRPATGTGSKVGKLTLKTTE | No |
| PRPF8 | Q6P2Q9 (2335) | 1772,<br>298,<br>427 | FSNQIIWFVDDTNVYRVTIHKTFEG<br>DINKIIIRQPIRTEYKIAFPYLYNN<br>VKNWYREHCPAGQPVKVRVSYQKLL | Partially |
| HSPA8 | NP_694881.1 (493) | 191,<br>216,<br>15 | AERNVLIFDLGGGTFDVSILTIEDG<br>IFEVKSTAGDTHLGGEDFDNRMVNH<br>YSCVGVFQHGKVEIIANDQGNRTTP | No |
| HSPA7 | P48741 (367) | 193,<br>127 | GKRNVLIFDLGGGTFDVSVLSIDAG<br>SKMKETAEAYLGQPVKHAVITVPTY | Partially |
| HSPA5 | P11021 (654) | 216,<br>504 | GEKNILVFDLGGGTFDVSLLTIDNG<br>DVNGILRVTAEDKGTGNKNKITITN | No |
| ACTR8 | Q9H981 (624) | 39,<br>240 | LQEQIQSNFIIVIHPGSTTLRIGRA<br>YNKQHVKELVNMILMKMGFSGIVVH | No |
| TAF3 | Q5VWG9 (929) | 760,<br>852 | RLTLRVGAGQDKIVISKVVPAPEAK<br>VSTYVIRDEWGNQIWICPGCNKPDD | Yes |
| TAF7 | Q15545(349) | 40,<br>13,<br>305 | KDRLTIELHPDGRHGIVRVDRVPLA<br>ESQFILRLPPEYASTVRRAVQSGHV<br>LIMKVENLALKNRFQAVLDELKQKE | Yes |
| NFRKB | Q6P4R8 (381) | 1169,<br>959,<br>635 | TTVVSTSQAGKLPTRITVPLSVISQ<br>SITTDAKGQTVLRITPDMMATLAKS<br>DPCVKYDIGRKLWIYLHRDRSEEEF | Yes |
| HSPA1L | P34931 (641) | 194,<br>484 | ERHVLIFDLGGGTFDVSILTIDDGI<br>ANGILNVTATDKSTGKVNKITITND | Partially |
| RPS6 | P62753 (249) | 1,<br>31,<br>133 | MKLNISFPATGCQKLIEVDDERKLR<br>RMATEVAADALGEEWKGYVVRISGG<br>LGPKRASRIRKLFNLSKEDDVRQYV | No |
| RPA1 | NP_001342050.1 (557) | 513,<br>73 | FRVRVKVETYNDESRIKATVMDVKP<br>SNCVCQIHRFIVNTLKDGRRVVILM | Yes |
| GLTSCR1 (BICRA) | Q9NZM4.2 (1560) | 1239,<br>1308,<br>441 | PPHLPTKLVIRHGGAGGSPSVTWAR<br>LKLKIKQEAGLSKVVHNTALDPVHQ<br>GALSKPMSVHLLNQGSSIVIPAQHM | Yes |
| SMARCA5 | O60264.1 (1052) | 626,<br>930,<br>496 | RLDSIVIQQGRL<br>APFHQLRISYGT<br>KEQGSRVLIFSQ | No |
| ZNF687 | NP_065883.1 (1237) | 328,<br>847 | LKVRIKTIKTSCGNITRTVTQVPSD<br>DQHLLPQRVSVFKCPSCPLLFAQKR | Partially |
| MED13 | Q9UHV7.3 (2174) | 494 | SQRLVISAPDSQVRFSNIRTNDVAK | Yes |
| CHD6 | NP_115597.3 (2715) | 988,<br>1725,<br>1762 | HTITIQSEGKGSTFAKASFVASGNR<br>STNTESRKDVITISISKDGNCQSGG<br>GSLEAGGVAQANIKNGKHLLMSISK | Yes |
| U2AF2 | P26368.4 (475) | 214,<br>412,<br>184 | FDGIIFQGQSLKIRRPHDYQPLPGM<br>VKSIEIPRPVDGVEVPGCGKIFVEF<br>NPVLAVQINQDKNFAFLEFRSVDET | Partially |
| ACTR1A | P61163.1 (376) | 177,<br>325 | PHSIMRIDIAGR<br>DVKIRISAPQER | Partially |
| INO80B | Q9C086.2 (356) | 64,<br>224 | PAKPQLKLKIKLGGQVLGTKSVPTF<br>RLQAARRAEEHKNQTIERLTKTAAT | Yes |

### Supporting Figures

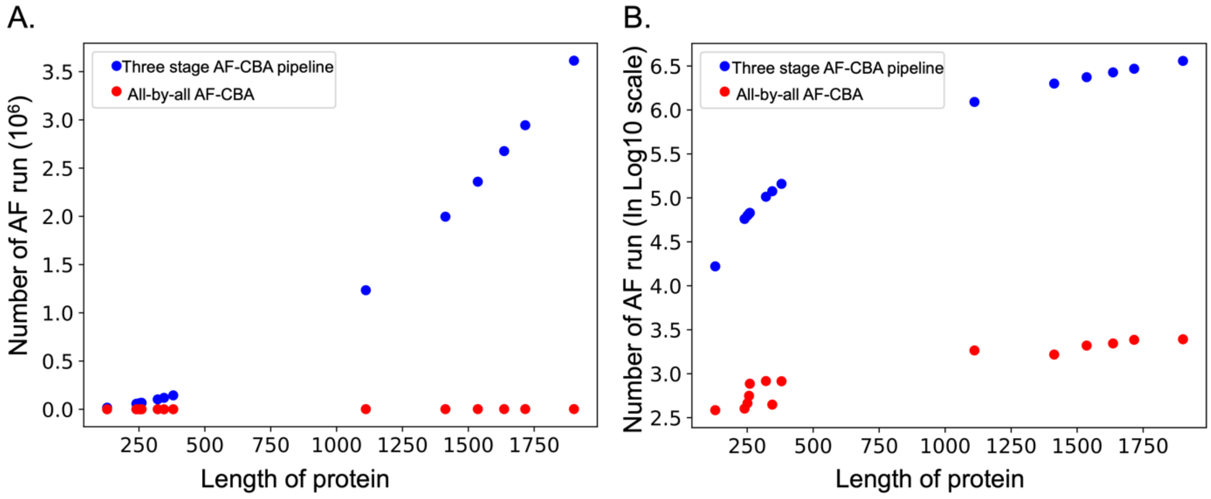


**Figure S1.** Computational cost comparison of two AF-CBA strategies shown in linear scale **(A)** and in logarithmic scale **(B).** We compare the AF-CBA pipeline proposed in this work vs a naïve all-by-all AF-CBA strategy.

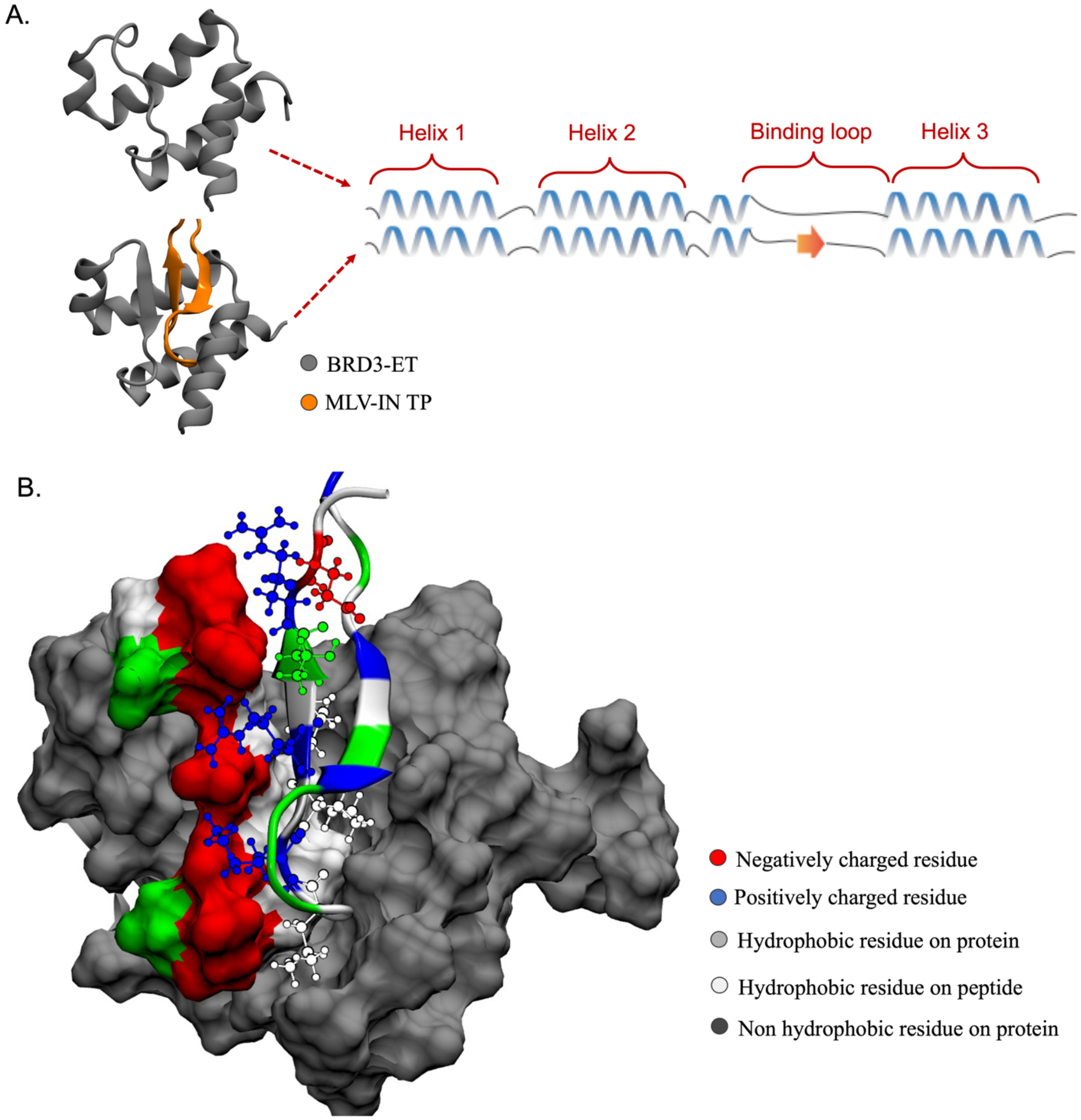


**Figure S2. (A)** Secondary structure elements in BRD3-ET in its unbound form and in complex with MLV-IN TP. **(B)** An alternating pattern of negative and hydrophobic residues in the ET receptor interacts with an alternating pattern of positive and hydrophobic residues in MLV-IN TP. This zipper like interaction is present in many of the known structures of ET in complex with peptides[9].

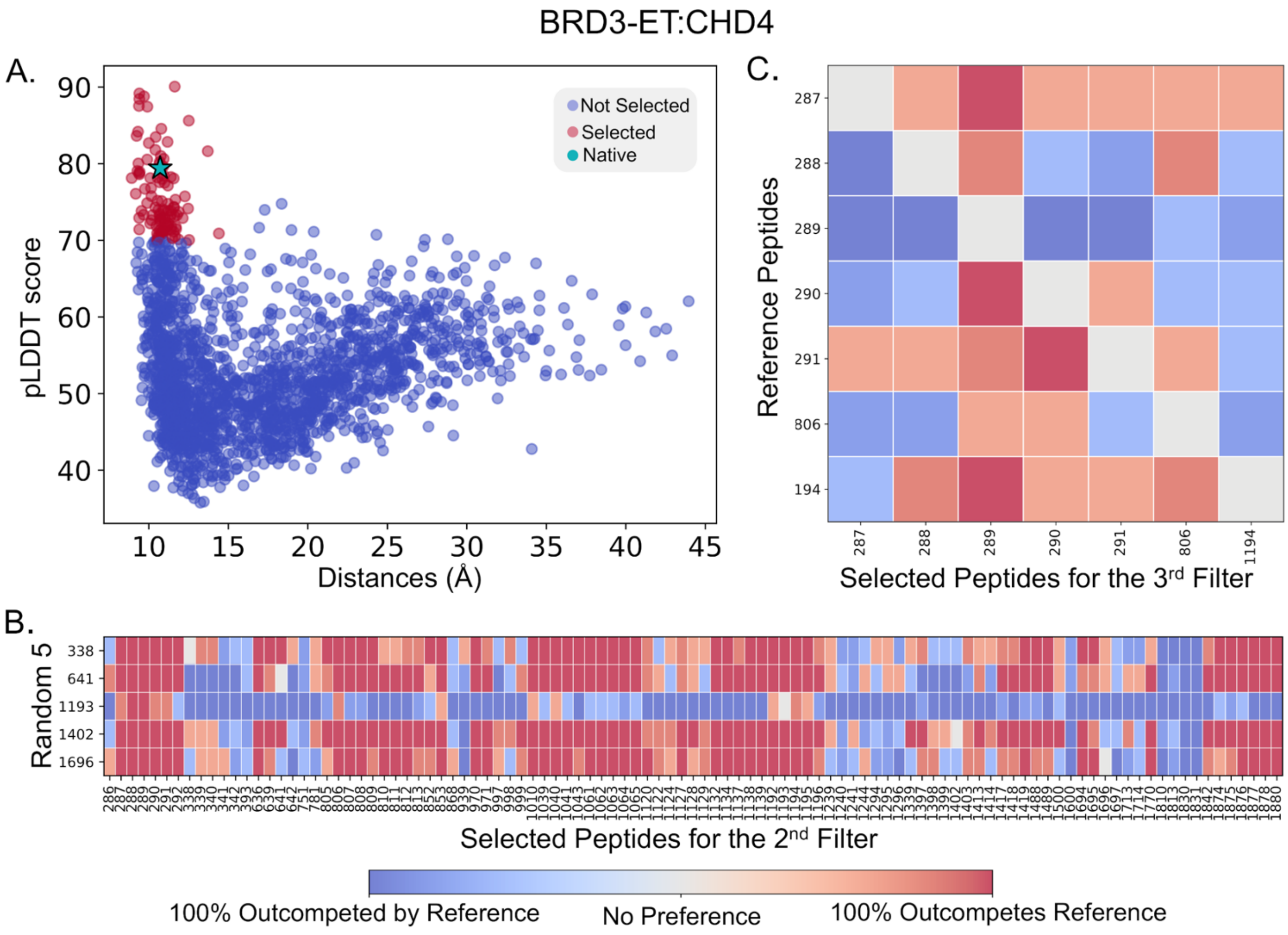


**Figure S3. Three step filtering results for BRD3-ET:CHD4**. **(A)** Filtering AF predictions for all possible epitopes against predicted pLDDT score (y axis) and average distance between all peptide residues to key residues in the receptor's binding site. Each dot is a prediction, with those in red advancing to the next stage, and the experimentally assigned peptide shown as a star. **(B)** AF-CBA between each peptide candidate in the remaining pool (each column) and each of five random peptides (each row). Columns with redder (see color bar) represent more successful binders and thus advance to the next filtering stage. **(C)** Remaining pool of peptides is evaluated in all-by-all AF-CBA experiment. Rows denotes peptides used as reference; columns identify each peptide to outcompete every other peptide used as reference. Columns with more red represent predicted peptides with higher binding affinity in the set.

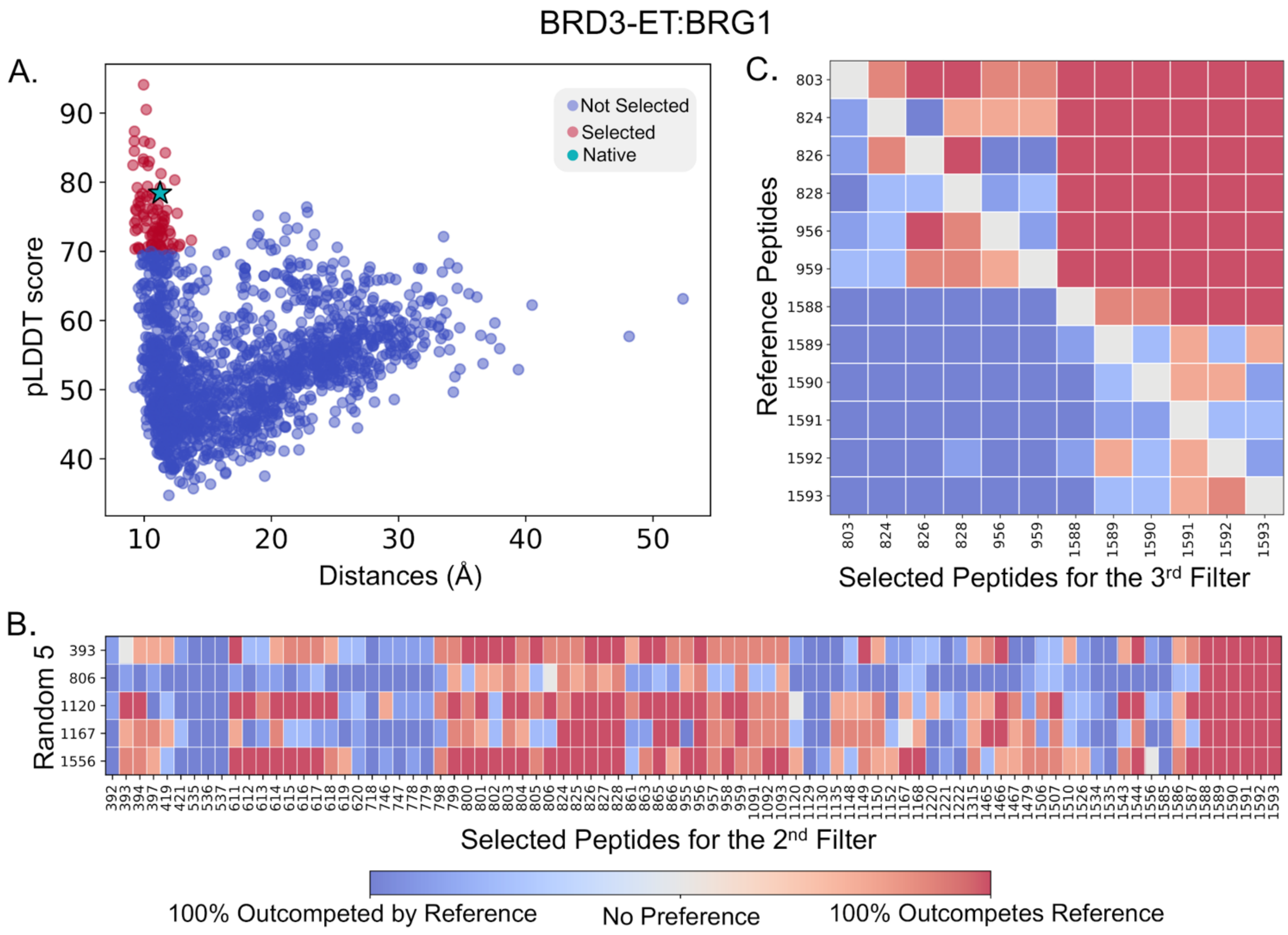


**Figure S4. Three step filtering results for BRD3-ET:BRG1**. **(A)** Filtering AF predictions for all possible epitopes against predicted pLDDT score (y axis) and average distance between all peptide residues to key residues in the receptor's binding site. Each dot is a prediction, with those in red advancing to the next stage, and the experimentally assigned peptide shown as a star. **(B)** AF-CBA between each peptide candidate in the remaining pool (each column) and each of five random peptides (each row). Columns with redder (see color bar) represent more successful binders and thus advance to the next filtering stage. **(C)** Remaining pool of peptides is evaluated in all-by-all AF-CBA experiment. Rows denotes peptides used as reference; columns identify each peptide to outcompete every other peptide used as reference. Columns with more red represent predicted peptides with higher binding affinity in the set.

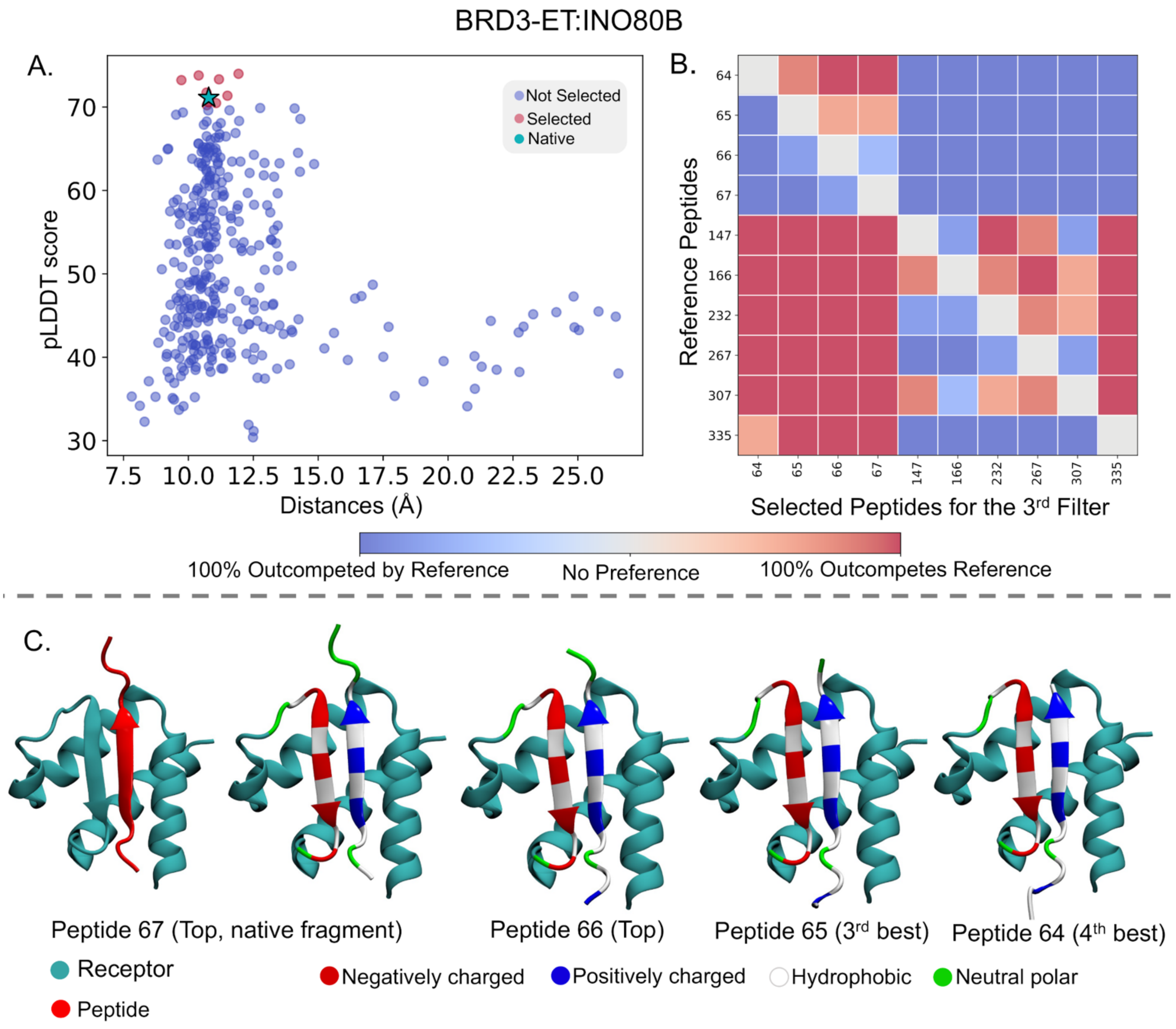


**Figure S5. Three step filtering results for BRD3-ET:INO80B**. **(A)** Filtering AF predictions for all possible epitopes against predicted pLDDT score (y axis) and average distance between all peptide residues to key residues in the receptor's binding site. Each dot is a prediction, with those in red advancing to the next stage, and the experimentally assigned peptide shown as a star **(B)** As <25 peptides passed the first stage, the remaining pool of peptides is evaluated in all-by-all AF-CBA experiment. Rows denotes peptides used as reference; columns identify each peptide to outcompete every other peptide used as reference. Columns with more red represent predicted peptides with higher binding affinity in the set. **(C)** Best prediction from AF-CBA along with 3 more neighbor and overlapping peptide fragments. These neighbor peptides perform similar to the best binder (Peptide 67) as their core binding residues are same with variable termini length.

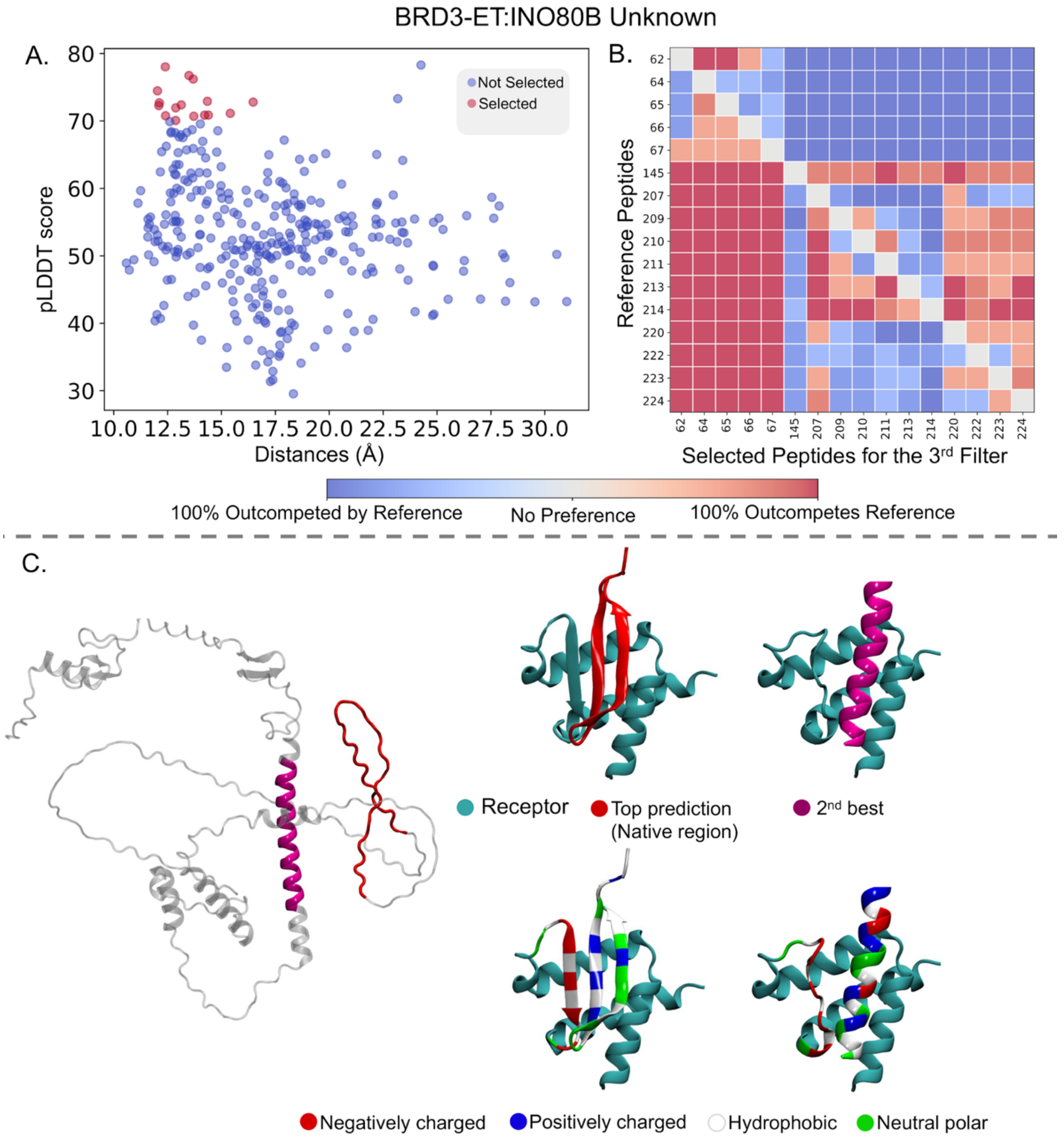


**Figure S6. Three step filtering results for BRD3-ET:INO80B (25 residues)**. **(A)** Filtering AF predictions for all possible epitopes against predicted pLDDT score (y axis) and average distance between all peptide residues to key residues in the receptor's binding site. Each dot is a prediction, with those in red advancing to the next stage, and the experimentally assigned peptide shown as a star **(B)** As <25 peptides passed the first stage, the remaining pool of peptides is evaluated in all-by-all AF-CBA experiment. Rows denotes peptides used as reference; columns identify each peptide to outcompete every other peptide used as reference. Columns with more red represent predicted peptides with higher binding affinity in the set. **(C)** AF predicted full length INO80B with two best binder regions highlighted (left) and complex structures of top two predicted epitopes with BRD3 (right). The best binder interacts through the same set of residues as the earlier 12-residues fragment.

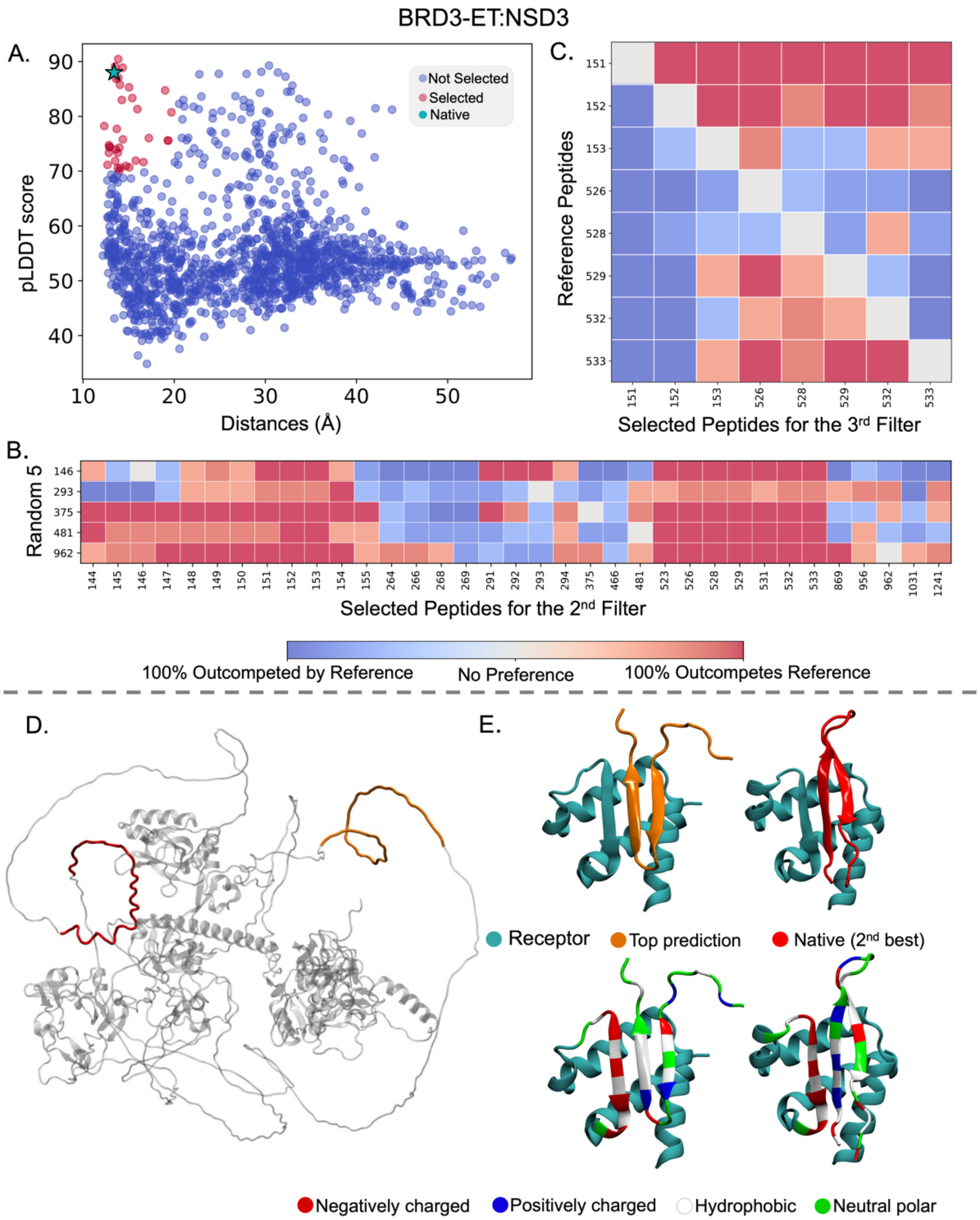


**Figure S7. Three step filtering results for BRD3-ET:NSD3**. **(A)** Filtering AF predictions for all possible epitopes against predicted pLDDT score (y axis) and average distance between all peptide residues to key residues in the receptor's binding site. Each dot is a prediction, with those in red advancing to the next stage, and the experimentally assigned peptide shown as a star. **(B)** AF-CBA between each peptide candidate in the remaining pool (each column) and each of five random peptides (each row). Columns with redder (see color bar) represent more successful binders and thus advance to the next filtering stage. **(C)** Remaining pool of peptides is evaluated in all-by-all AF-CBA experiment. Rows denotes peptides used as reference; columns identify each peptide to outcompete every other peptide used as reference. Columns with more red represent predicted peptides with higher binding affinity in the set. **(D)** AF predicted full length NSD3 with two best binder regions highlighted (left) and complex structures of top two predicted epitopes with BRD3 (right). The experimentally known binder (canonical) comes out as the 2nd best binder.

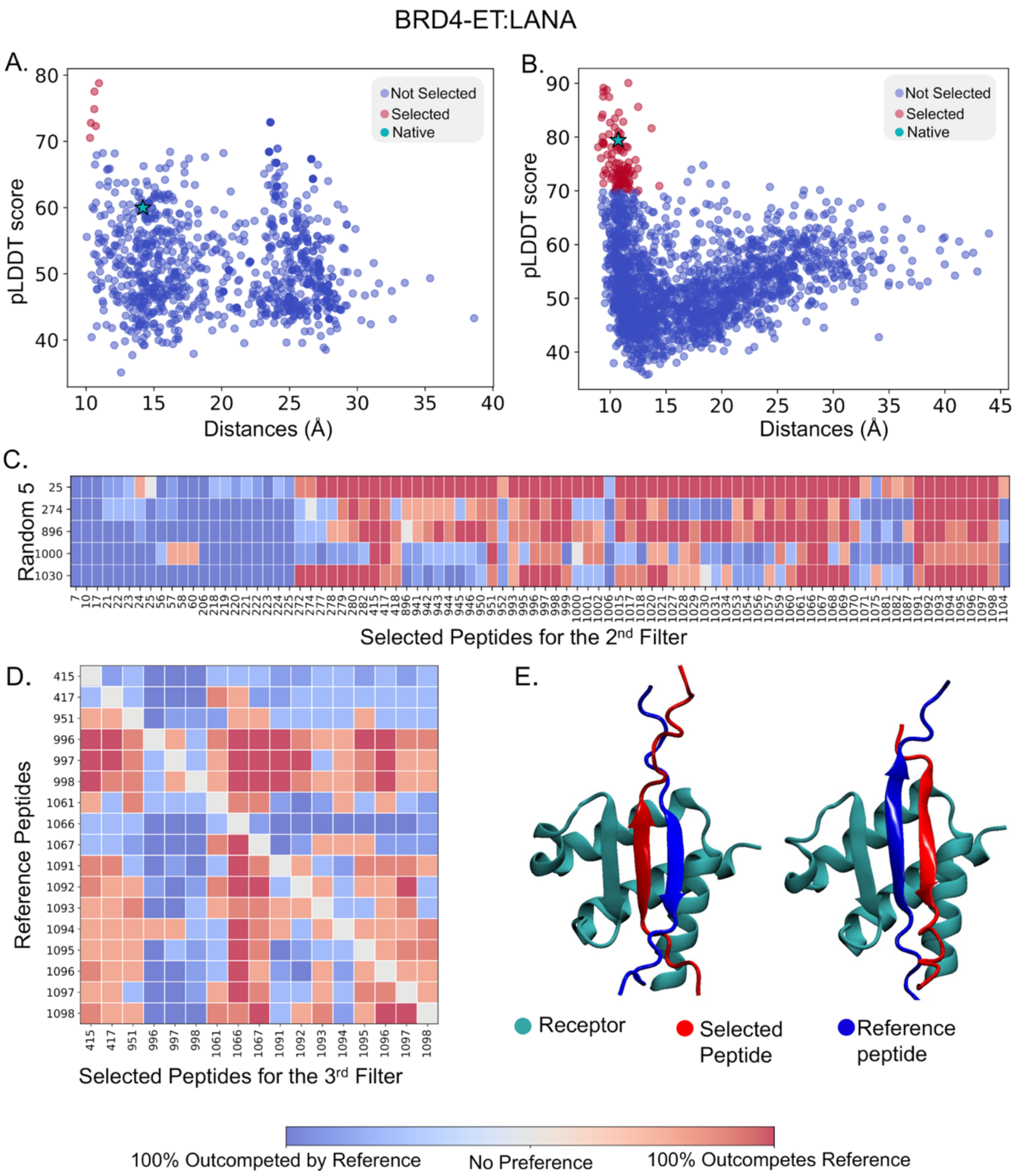


**Figure S8. Three step filtering results for BRD4-ET:LANA**. **(A)** Filtering AF predictions for all possible epitopes against predicted pLDDT score (y axis) and average distance between all peptide residues to key residues in the receptor's binding site. Each dot is a prediction, with those in red advancing to the next stage, and the experimentally assigned peptide shown as a star. Here native binder fragment is not selected when pLDDT cutoff is 70. **(B)** Relaxing cutoff to 55 can select the native binder. **(C)** AF-CBA between each peptide candidate in the remaining pool (each column) and each of five random peptides (each row). Columns with redder (see color bar) represent more successful binders and thus advance to the next filtering stage. **(D)** Remaining pool of peptides is evaluated in all-by-all AF-CBA experiment. Rows denotes peptides used as reference; columns identify each peptide to outcompete every other peptide used as reference. Columns with more red represent predicted peptides with higher binding affinity in the set. **(D)** Co-binding of two competing peptides at the 3rd filtering stage.

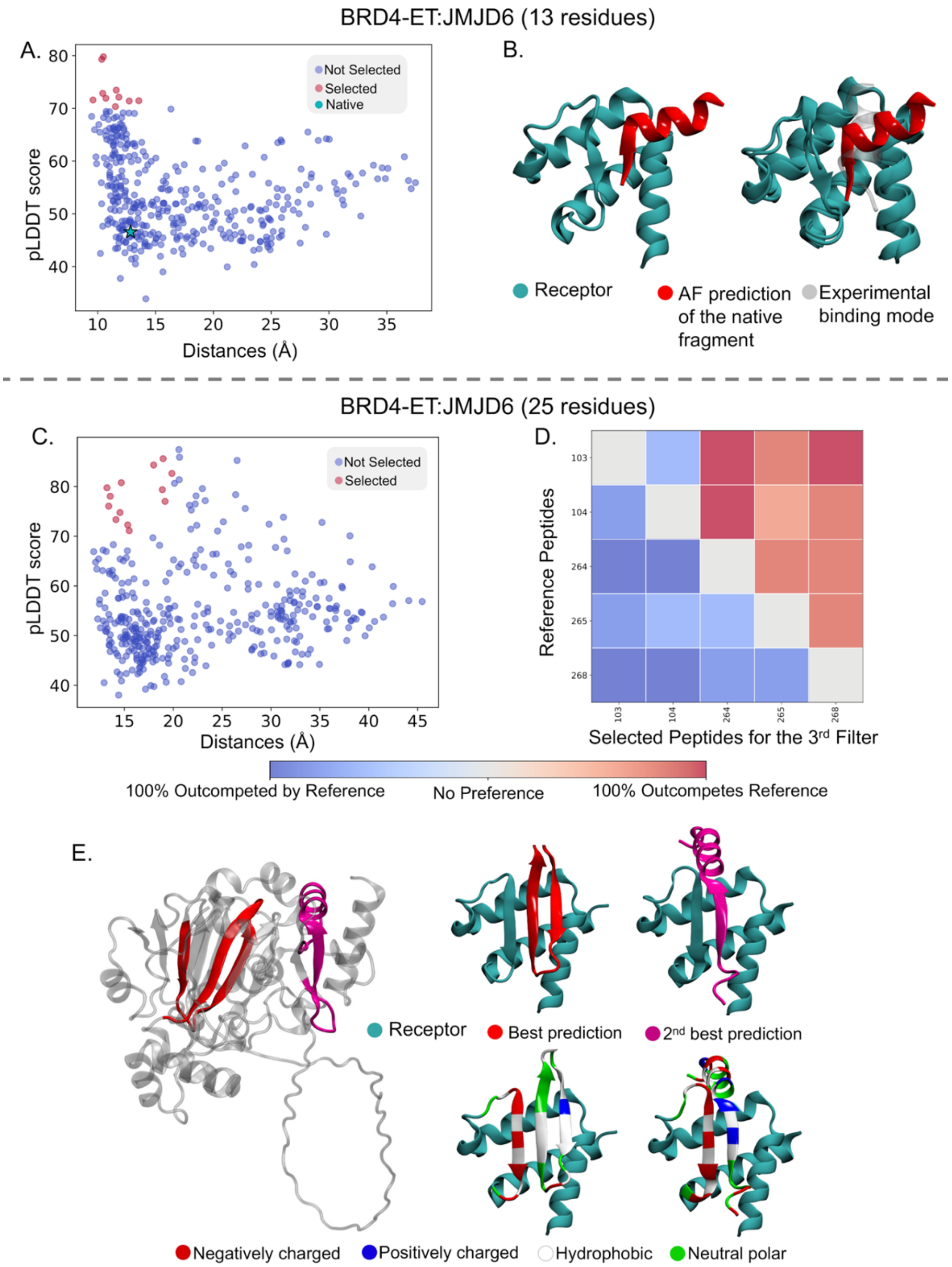


**Figure S9. Three step filtering results for BRD4-ET:JMJD6. (A)** Filtering AF predictions for all possible epitopes against predicted pLDDT score (y axis) and average distance between all peptide residues to key residues in the receptor's binding site. Each dot is a prediction, with those in red advancing to the next stage, and the experimentally assigned peptide shown as a star. The native binder is not selected. **(B)** AF fails to predict the native binder epitope. **(C)** AF binding of peptides fragments with the length of 25 residues. **(D)** Remaining pool of peptides is evaluated in all-by-all AF-CBA experiment. Rows denotes peptides used as reference; columns identify each peptide to outcompete every other peptide used as reference. Columns with more red represent predicted peptides with higher binding affinity in the set. **(E)** AF predicted full length JMJD6 with two best binder regions highlighted (left) and complex structures of top two predicted epitopes with BRD4 (right).

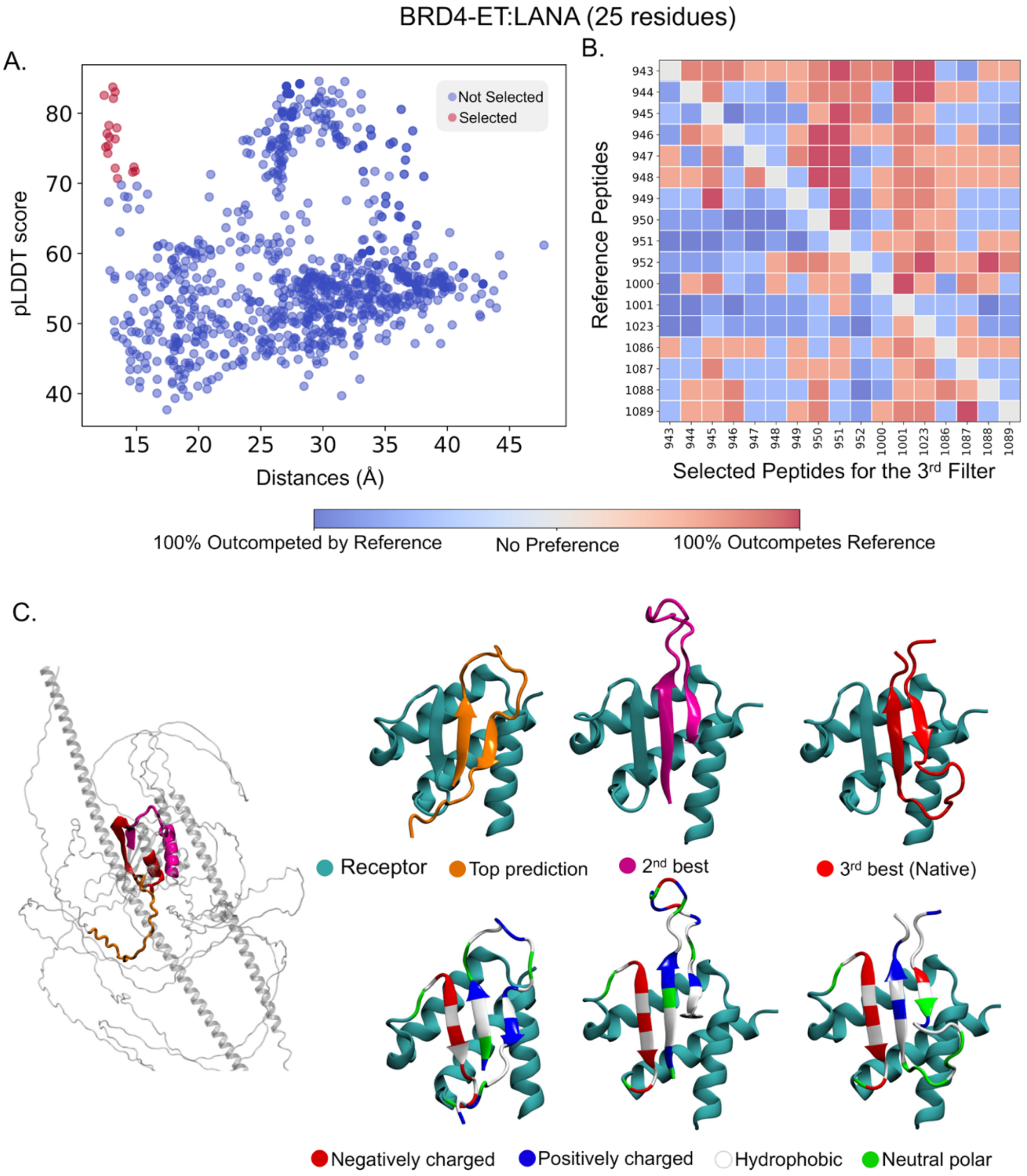


**Figure S10. Three step filtering results for BRD4-ET:LANA (25 residue)**. **(A)** Filtering AF predictions for all possible epitopes against predicted pLDDT score (y axis) and average distance between all peptide residues to key residues in the receptor's binding site. Each dot is a prediction, with those in red advancing to the next stage, and the experimentally assigned peptide shown as a star **(B)** Remaining pool of peptides is evaluated in all-by-all AF-CBA experiment. Rows denotes peptides used as reference; columns identify each peptide to outcompete every other peptide used as reference. Columns with more red represent predicted peptides with higher binding affinity in the set. **(C)** AF predicted full length LANA with three best binder regions highlighted (left) and complex structures of top three predicted epitopes with BRD4 (right). The experimentally known binder (canonical) comes out as the 3$^{rd}$ best binder as a hairpin due to extending the length of the fragment.

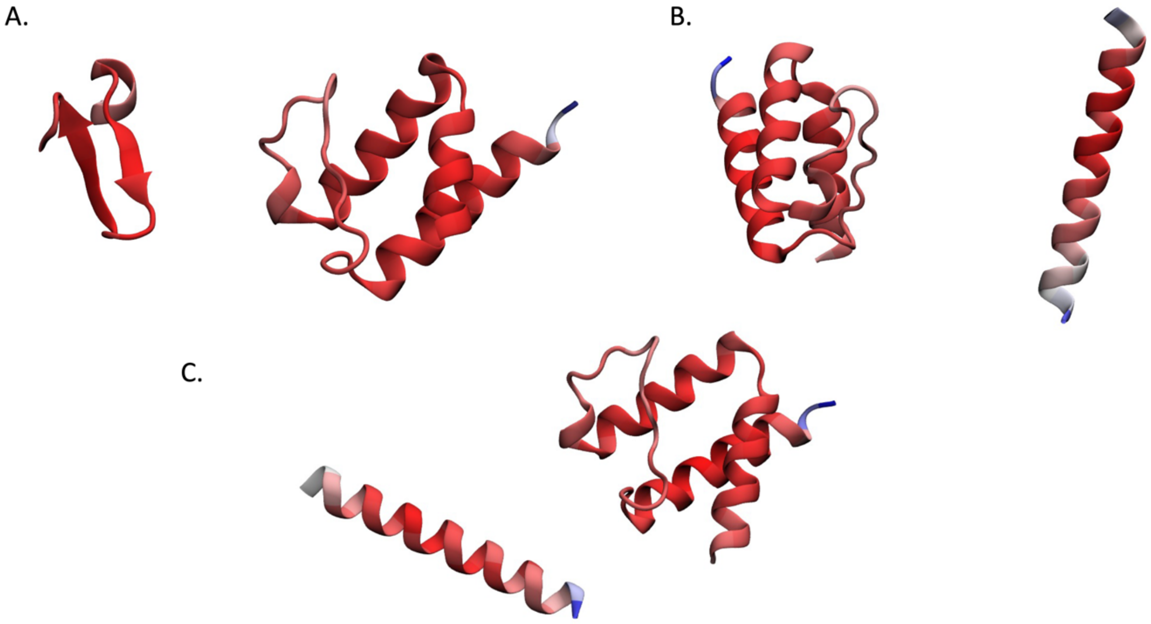


**Figure S11**. Example of AF predicted complexes with high pLDDT score, but the peptides are not bound to the receptors for TP (Fragment 1350) **(A)**, NSD3 (Fragment 1040) **(B),** and LANA (Fragment 796) **(C).** AF predicted per residue pLDDT score is represented with a red to white to blue colors scale ranging pLDDT values from 100 to 50 to 0.

1NLN

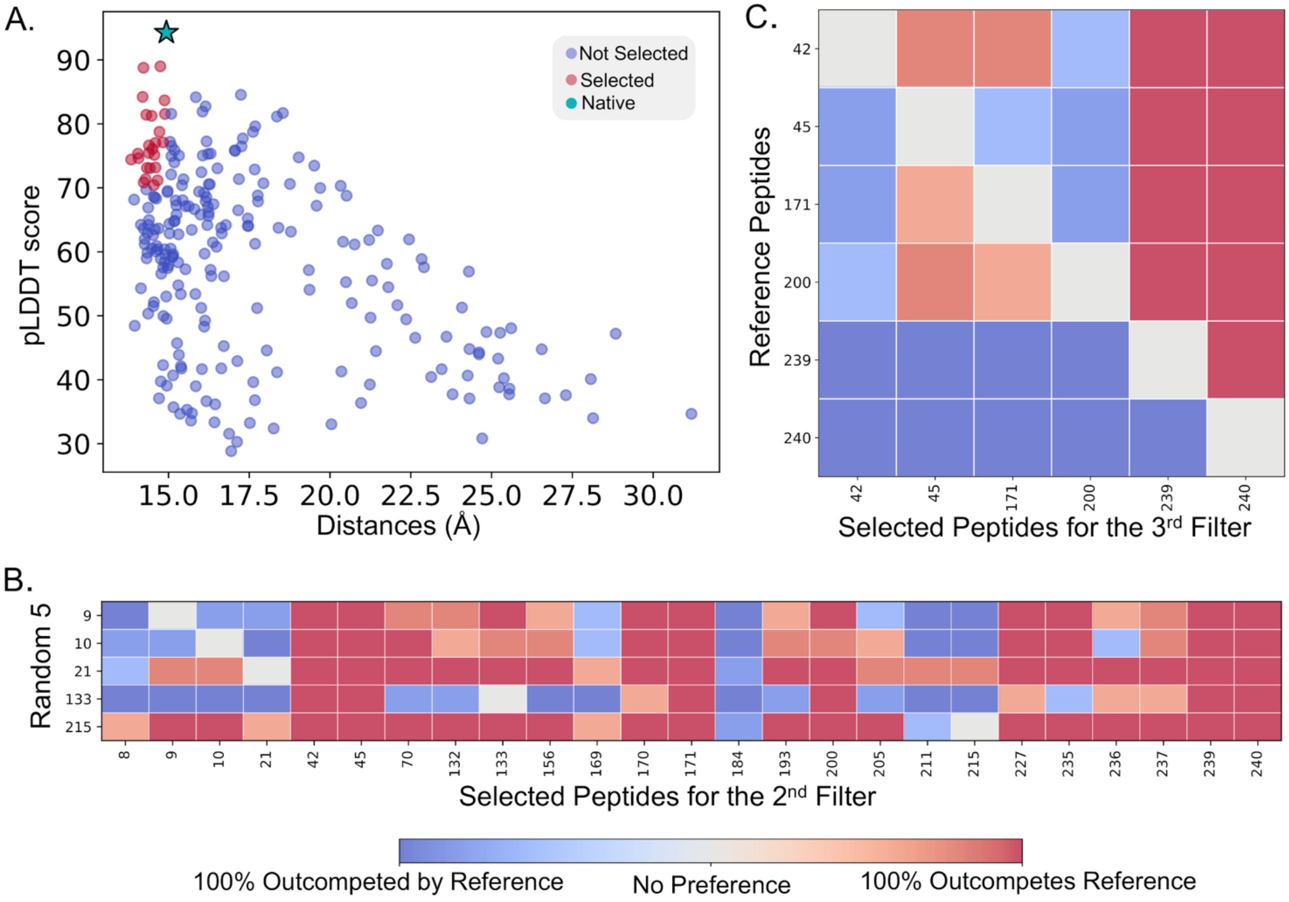


**Figure S12. Three step filtering results for 1NLN**. **(A)** Filtering AF predictions for all possible epitopes against predicted pLDDT score (y axis) and average distance between all peptide residues to key residues in the receptor's binding site. Each dot is a prediction, with those in red advancing to the next stage, and the experimentally assigned peptide shown as a star. **(B)** AF-CBA between each peptide candidate in the remaining pool (each column) and each of five random peptides (each row). Columns with redder (see color bar) represent more successful binders and thus advance to the next filtering stage. **(C)** Remaining pool of peptides is evaluated in all-by-all AF-CBA experiment. Rows denotes peptides used as reference; columns identify each peptide to outcompete every other peptide used as reference. Columns with more red represent predicted peptides with higher binding affinity in the set.

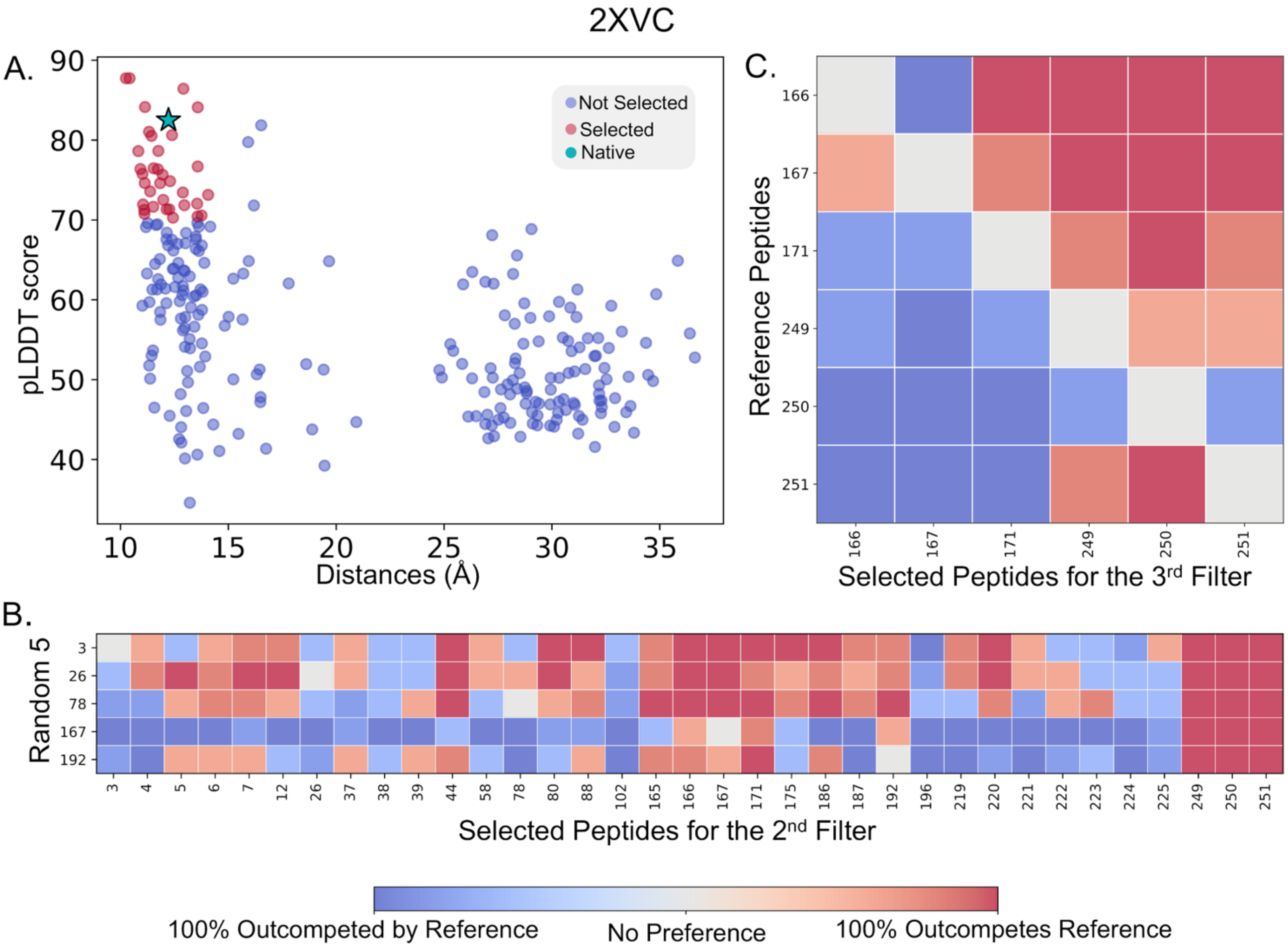


**Figure S13. Three step filtering results for 2XVC**. **(A)** Filtering AF predictions for all possible epitopes against predicted pLDDT score (y axis) and average distance between all peptide residues to key residues in the receptor's binding site. Each dot is a prediction, with those in red advancing to the next stage, and the experimentally assigned peptide shown as a star. **(B)** AF-CBA between each peptide candidate in the remaining pool (each column) and each of five random peptides (each row). Columns with redder (see color bar) represent more successful binders and thus advance to the next filtering stage. **(C)** Remaining pool of peptides is evaluated in all-by-all AF-CBA experiment. Rows denotes peptides used as reference; columns identify each peptide to outcompete every other peptide used as reference. Columns with more red represent predicted peptides with higher binding affinity in the set.

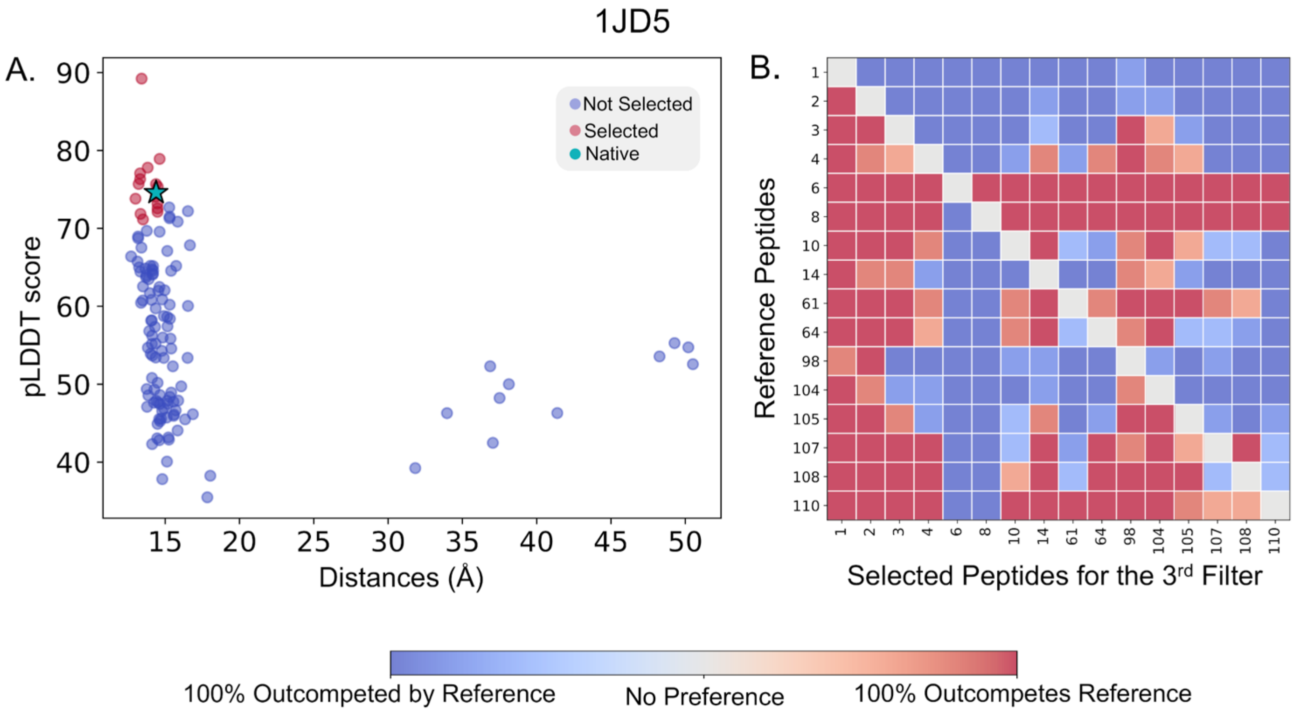


**Figure S14. Three step filtering results for 1JD5**. **(A)** Filtering AF predictions for all possible epitopes against predicted pLDDT score (y axis) and average distance between all peptide residues to key residues in the receptor's binding site. Each dot is a prediction, with those in red advancing to the next stage, and the experimentally assigned peptide shown as a star **(B)** As <25 peptides passed the first stage, the remaining pool of peptides is evaluated in all-by-all AF-CBA experiment. Rows denotes peptides used as reference; columns identify each peptide to outcompete every other peptide used as reference. Columns with more red represent predicted peptides with higher binding affinity in the set.

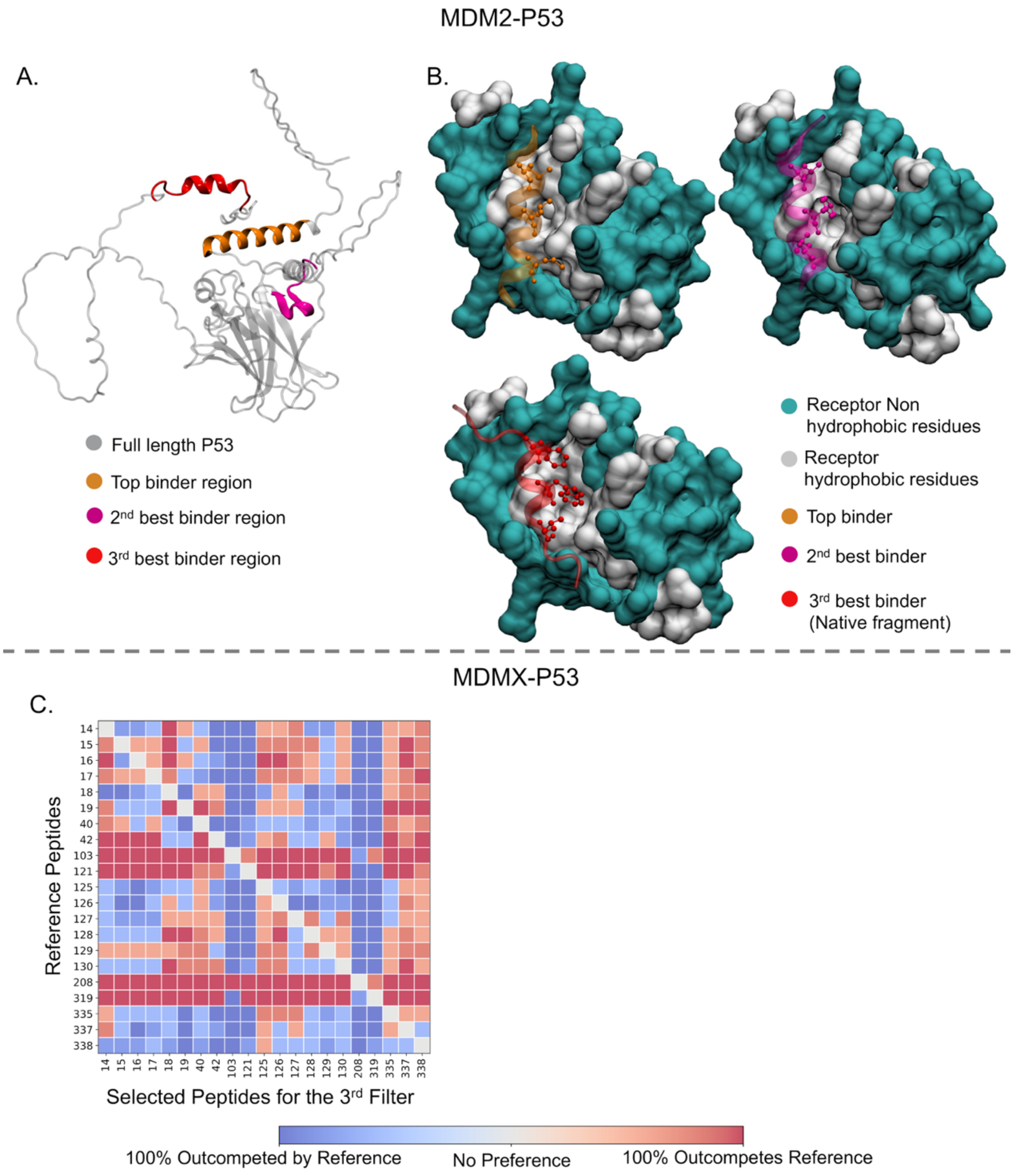


**Figure S15. Three step filtering results for MDM2-P53 and MDMX-P53. (A)** AF predicted full length P53 with three best binder regions highlighted. **(B)** Complex structures of top three predicted epitopes with. The experimentally known binder (canonical) comes out as the 3rd best binder. The other two best binders also bind anchoring 3 hydrophobic residues to the hydrophobic pocket of MDM2. **(C)** The selected peptides for the 3rd filter from MDM2 binding (See Figure 4) are tested against MDMX. Tested pool of peptides is evaluated in all-by-all AF-CBA experiment. Rows denotes peptides used as reference; columns identify each peptide to outcompete every other peptide used as reference. Columns with more red represent predicted peptides with higher binding affinity in the set.

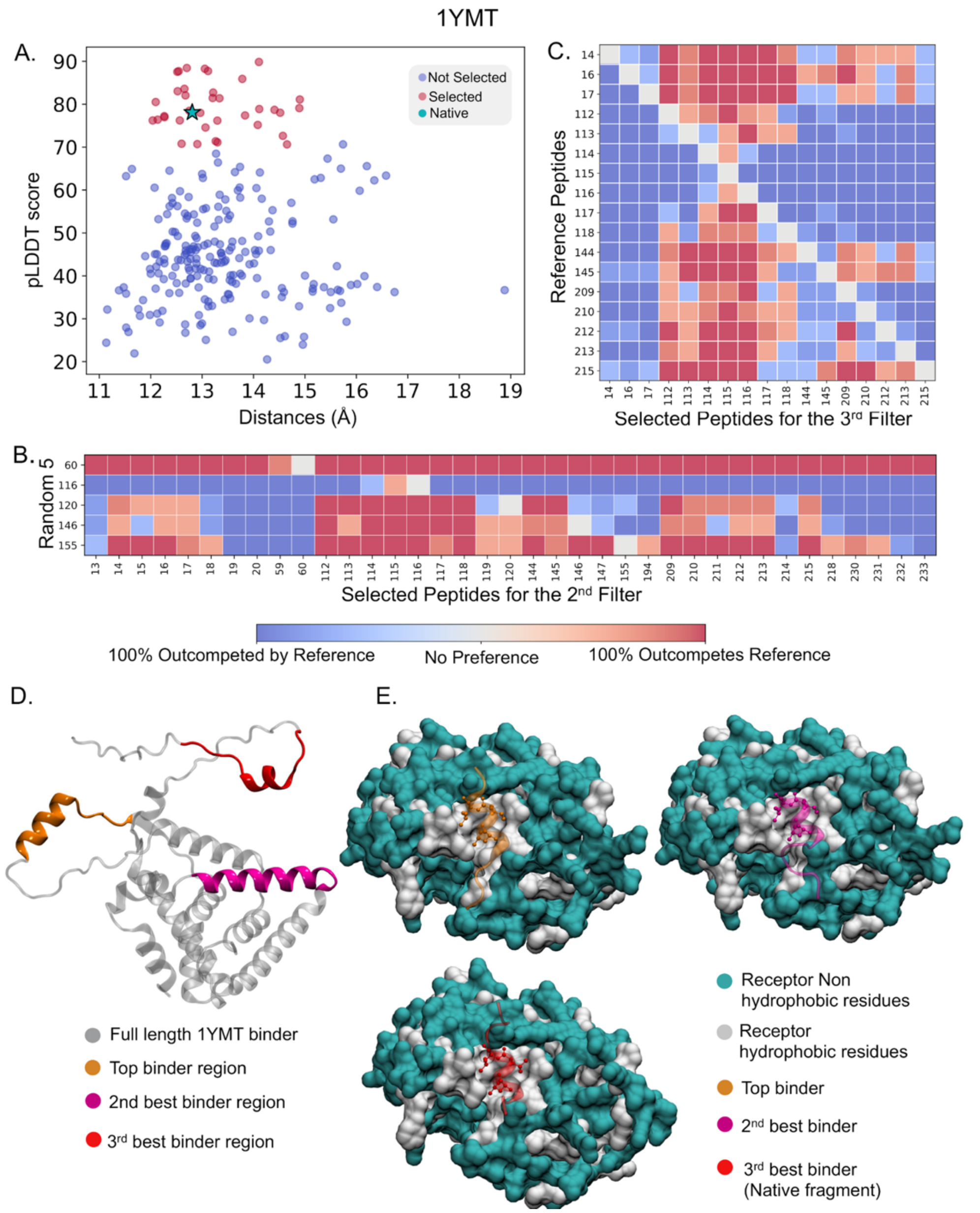


**Figure S16. Three step filtering results for 1YMT**. **(A)** Filtering AF predictions for all possible epitopes against predicted pLDDT score (y axis) and average distance between all peptide residues to key residues in the receptor's binding site. Each dot is a prediction, with those in red advancing to the next stage, and the experimentally assigned peptide shown as a star. **(B)** AF-CBA between each peptide candidate in the remaining pool (each column) and each of five random peptides (each row). Columns with redder (see color bar) represent more successful binders and thus advance to the next filtering stage. **(C)** Remaining pool of peptides is evaluated in all-by-all AF-CBA experiment. Rows denotes peptides used as reference; columns identify each peptide to outcompete every other peptide used as reference. Columns with more red represent predicted peptides with higher binding affinity in the set. **(D)** AF predicted full length binder with three best binder regions highlighted. **(F)** Complex structures of top three predicted epitopes with the receptor. The experimentally known binder (canonical) comes out as the 3rd best binder. The other two best binders also bind anchoring 3 hydrophobic residues to the hydrophobic pocket of the receptor.

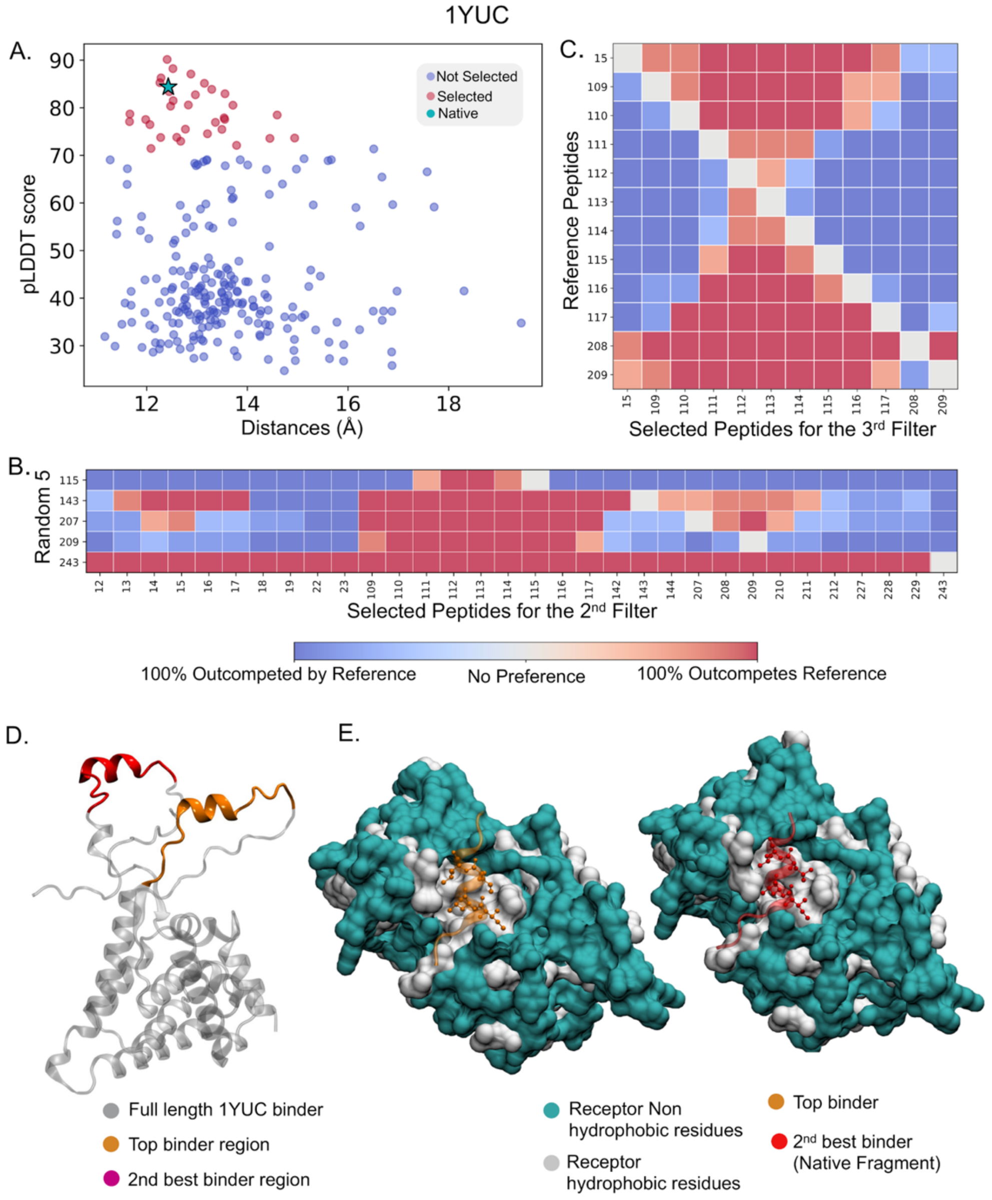


**Figure S17. Three step filtering results for 1YUC**. **(A)** Filtering AF predictions for all possible epitopes against predicted pLDDT score (y axis) and average distance between all peptide residues to key residues in the receptor's binding site. Each dot is a prediction, with those in red advancing to the next stage, and the experimentally assigned peptide shown as a star. **(B)** AF-CBA between each peptide candidate in the remaining pool (each column) and each of five random peptides (each row). Columns with redder (see color bar) represent more successful binders and thus advance to the next filtering stage. **(C)** Remaining pool of peptides is evaluated in all-by-all AF-CBA experiment. Rows denotes peptides used as reference; columns identify each peptide to outcompete every other peptide used as reference. Columns with more red represent predicted peptides with higher binding affinity in the set. **(D)** AF predicted full length binder with two best binder regions highlighted. **(F)** Complex structures of top two predicted epitopes with the receptor. The experimentally known binder (canonical) comes out as the 2nd best binder. The best binders also bind anchoring 3 hydrophobic residues to the hydrophobic pocket of the receptor.

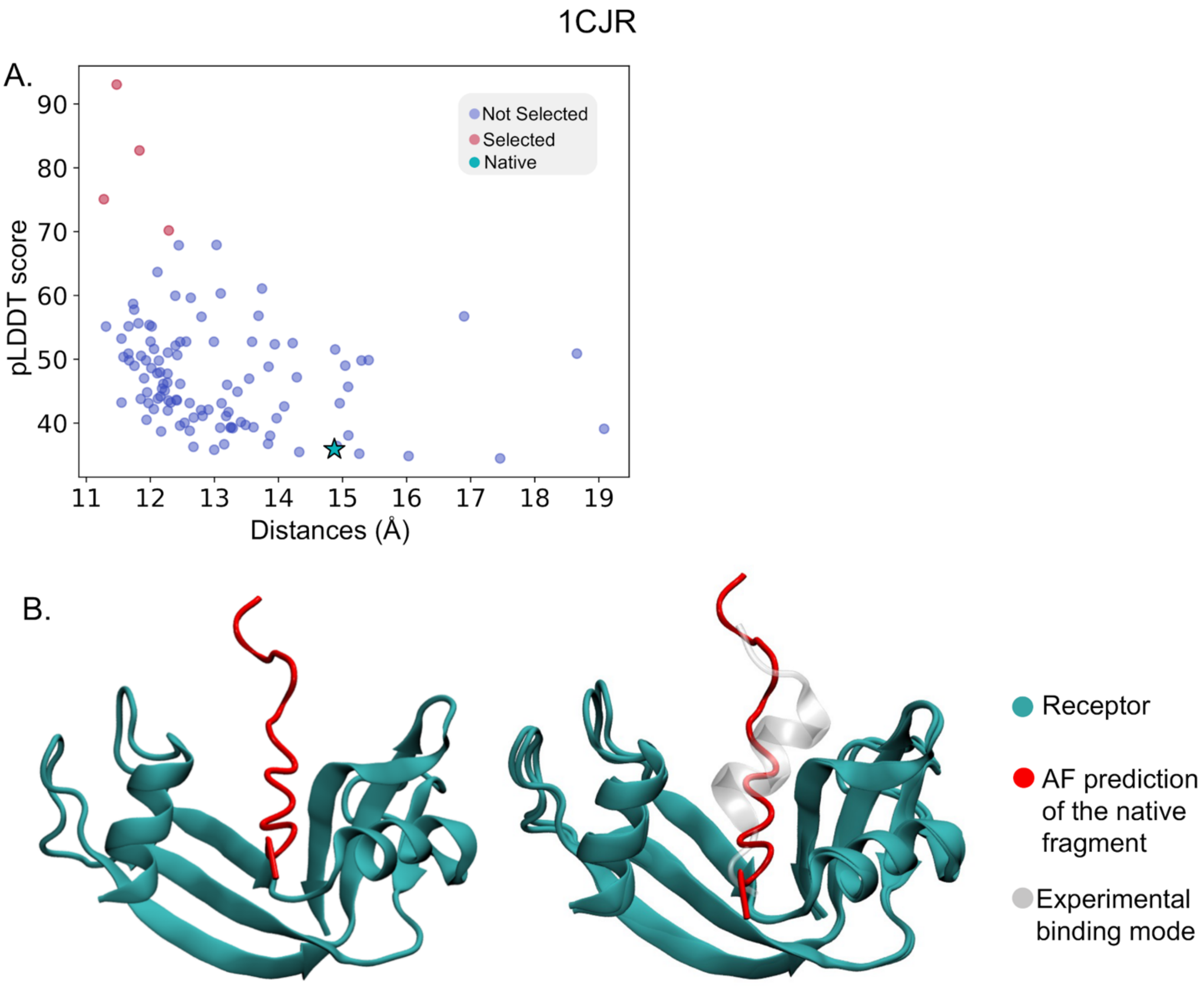


**Figure S18. Three step filtering results for 1CJR. (A)** Filtering AF predictions for all possible epitopes against predicted pLDDT score (y axis) and average distance between all peptide residues to key residues in the receptor's binding site. Each dot is a prediction, with those in red advancing to the next stage, and the experimentally assigned peptide shown as a star. The native binder is not selected. **(B)** AF fails to predict the native binder epitope. AF predicts an unstructured coil instead of a helix.

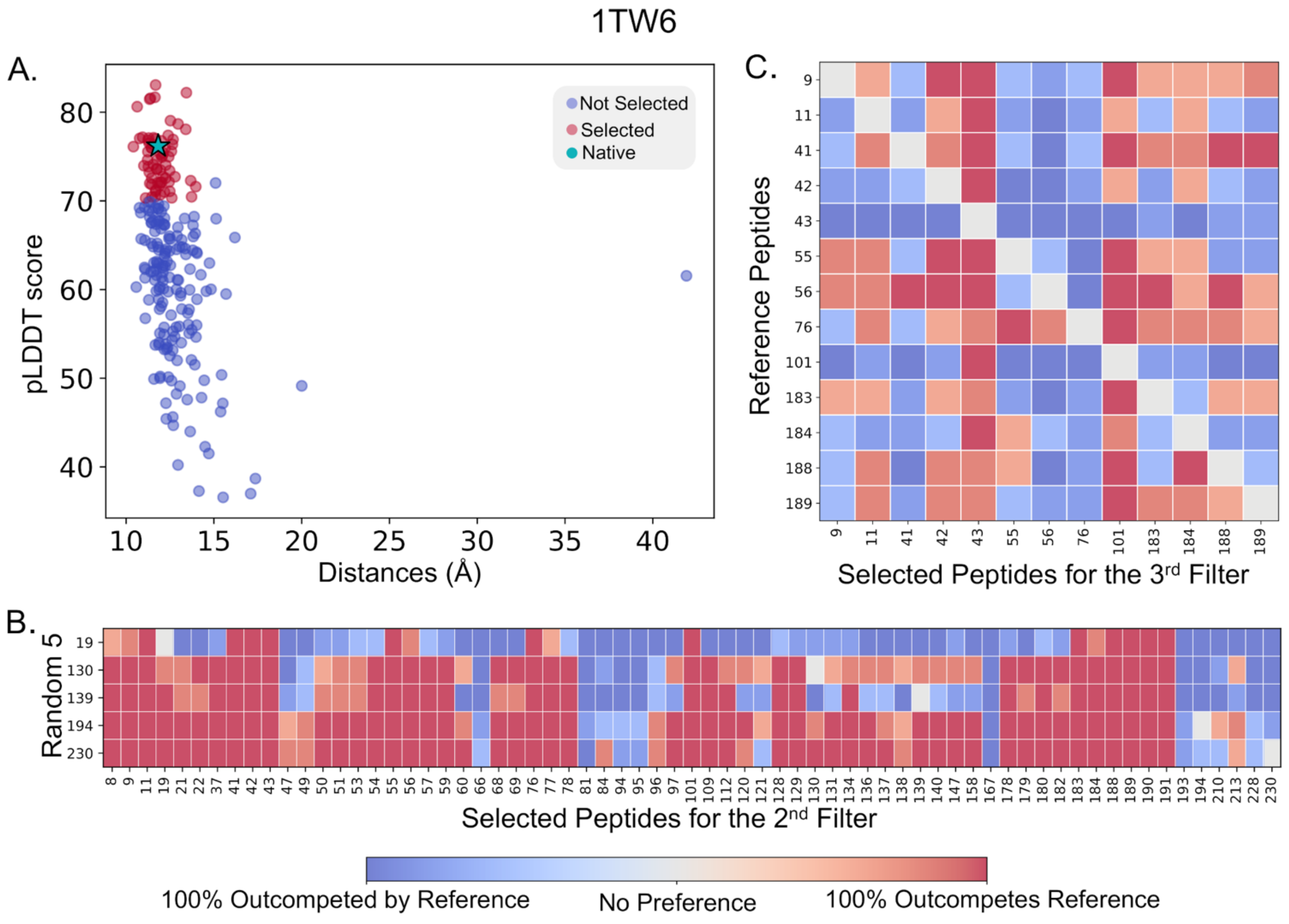


**Figure S19. Three step filtering results for 1TW6**. **(A)** Filtering AF predictions for all possible epitopes against predicted pLDDT score (y axis) and average distance between all peptide residues to key residues in the receptor's binding site. Each dot is a prediction, with those in red advancing to the next stage, and the experimentally assigned peptide shown as a star. **(B)** AF-CBA between each peptide candidate in the remaining pool (each column) and each of five random peptides (each row). Columns with redder (see color bar) represent more successful binders and thus advance to the next filtering stage. **(C)** Remaining pool of peptides is evaluated in all-by-all AF-CBA experiment. Rows denotes peptides used as reference; columns identify each peptide to outcompete every other peptide used as reference. Columns with more red represent predicted peptides with higher binding affinity in the set.

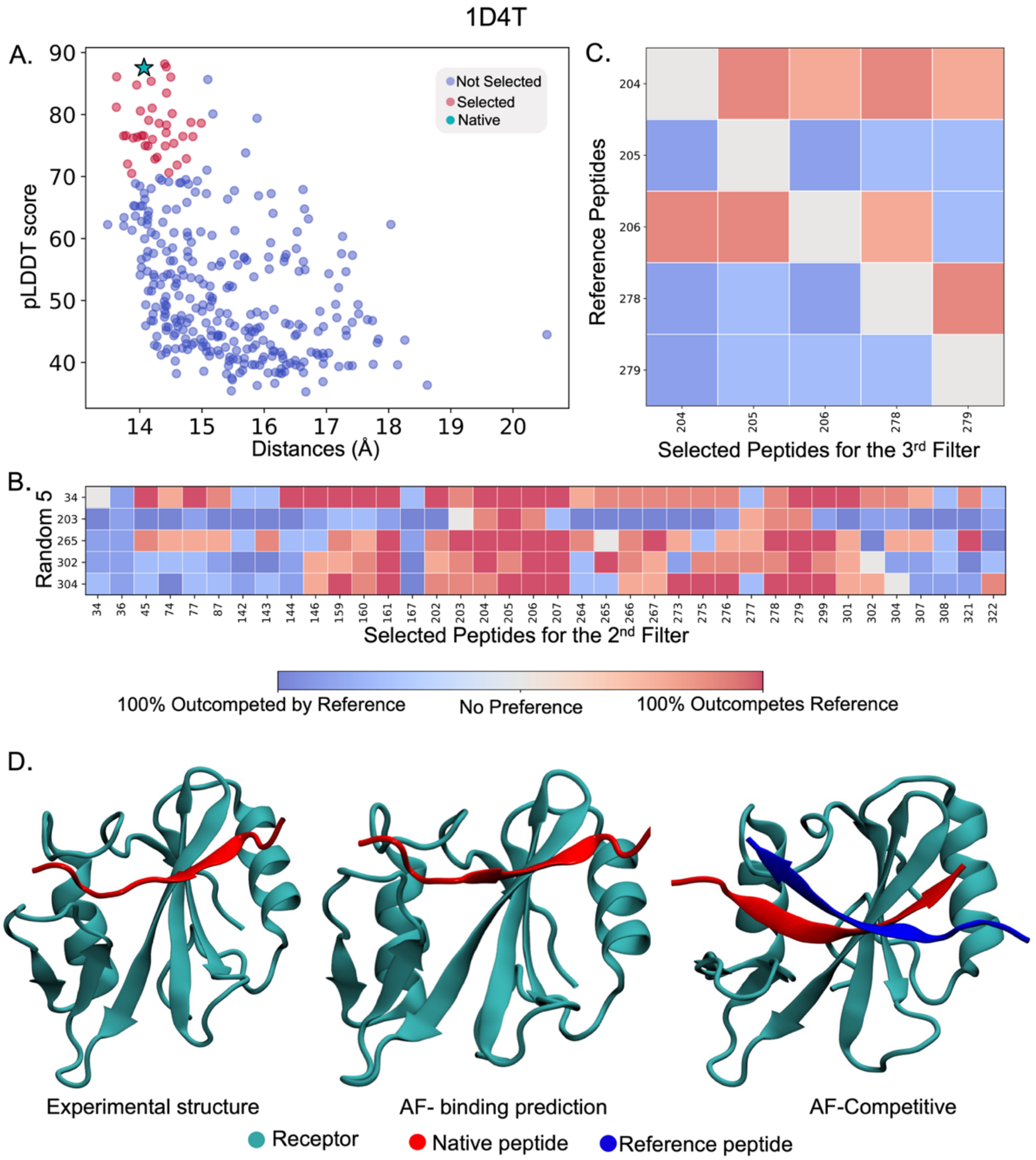


**Figure S20. Three step filtering results for 1D4T**. **(A)** Filtering AF predictions for all possible epitopes against predicted pLDDT score (y axis) and average distance between all peptide residues to key residues in the receptor's binding site. Each dot is a prediction, with those in red advancing to the next stage, and the experimentally assigned peptide shown as a star. **(B)** AF-CBA between each peptide candidate in the remaining pool (each column) and each of five random peptides (each row). Columns with redder (see color bar) represent more successful binders and thus advance to the next filtering stage. **(C)** Remaining pool of peptides is evaluated in all-by-all AF-CBA experiment. Rows denotes peptides used as reference; columns identify each peptide to outcompete every other peptide used as reference. Columns with more red represent predicted peptides with higher binding affinity in the set. **(D)** The native binding epitope co-binds with other competing peptides at the competitive binding stage in a flipped orientation suggesting the unreliability of the prediction.

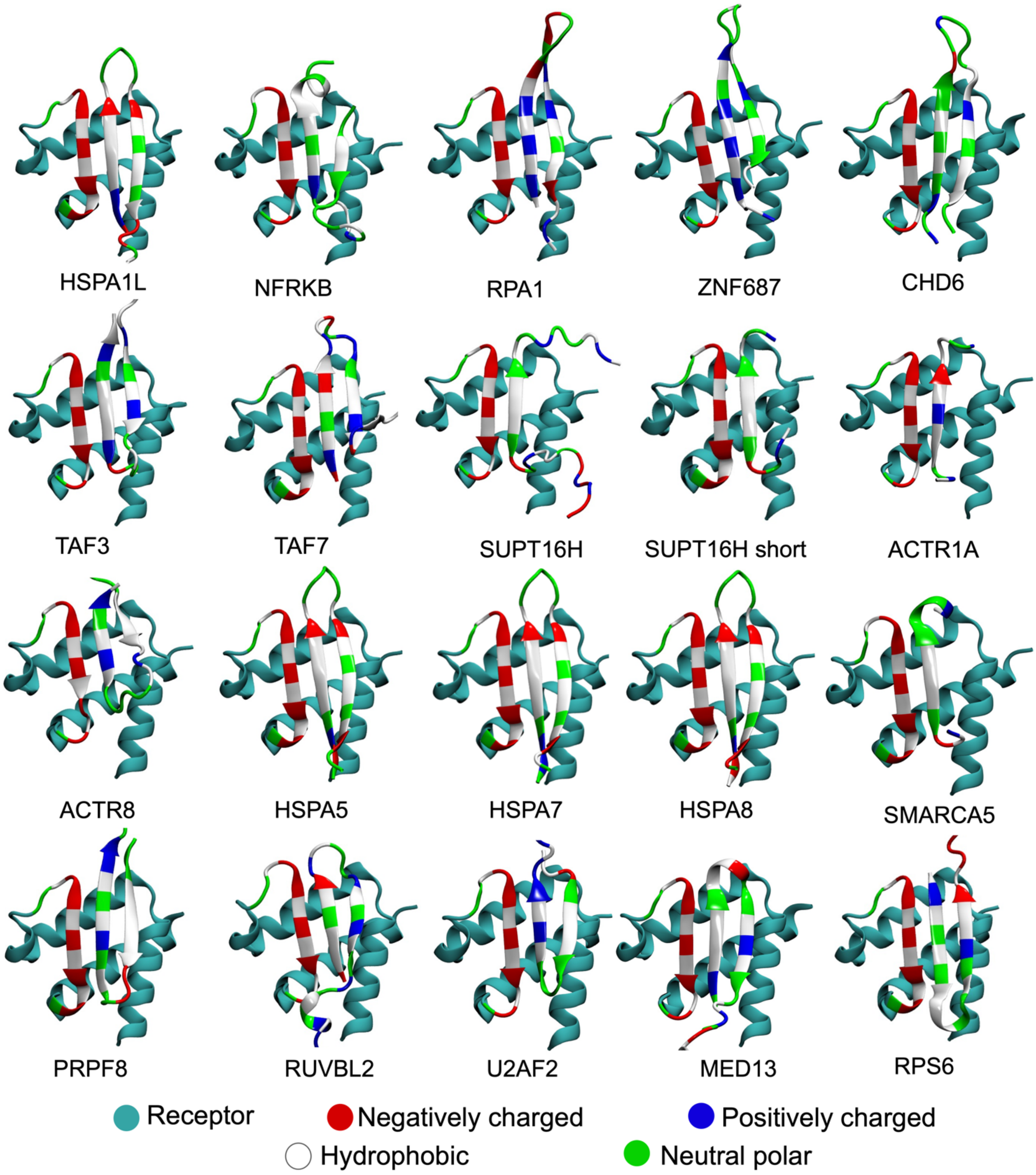


**Figure S21.** AF predicted structures of BRD3-ET complexes with the best predicted peptide epitopes derived from high confidence pull down hits. The peptide and binding region of receptors are shown in residue type color to rapidly visualize canonical vs non-canonical bioinformatic motifs (alternate charge/hydrophobic patterns). The receptor is shown in cyan.

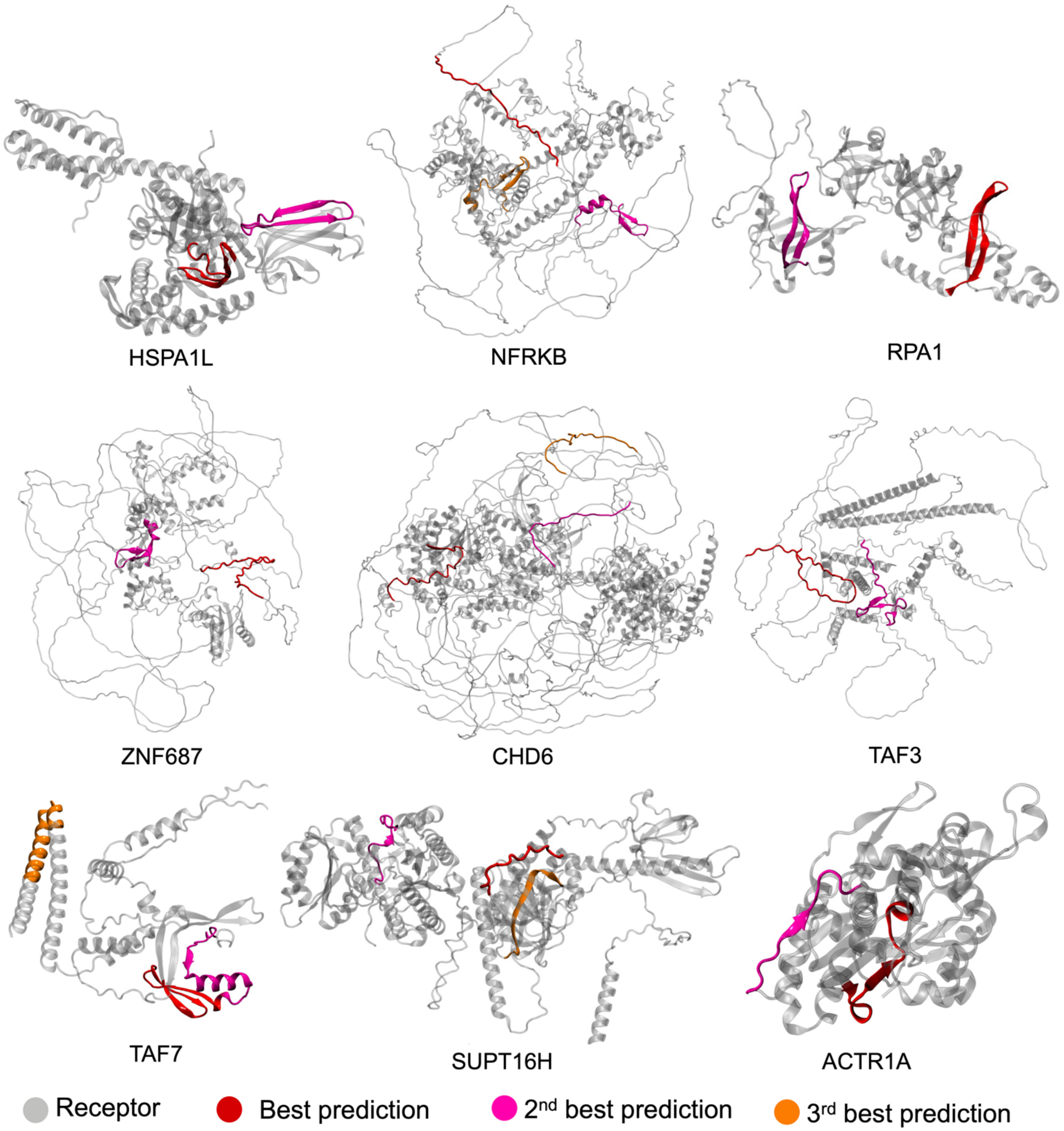


**Figure S22.** Full-length AF predicted structures of protein sequences derived from high-confidence pull-down hits. Top predicted epitope regions are shown red, magenta, and orange color. Rest of the proteins are shown in silver. AF regions that are not predicted have been observed to often correspond to intrinsically disordered regions in the protein.

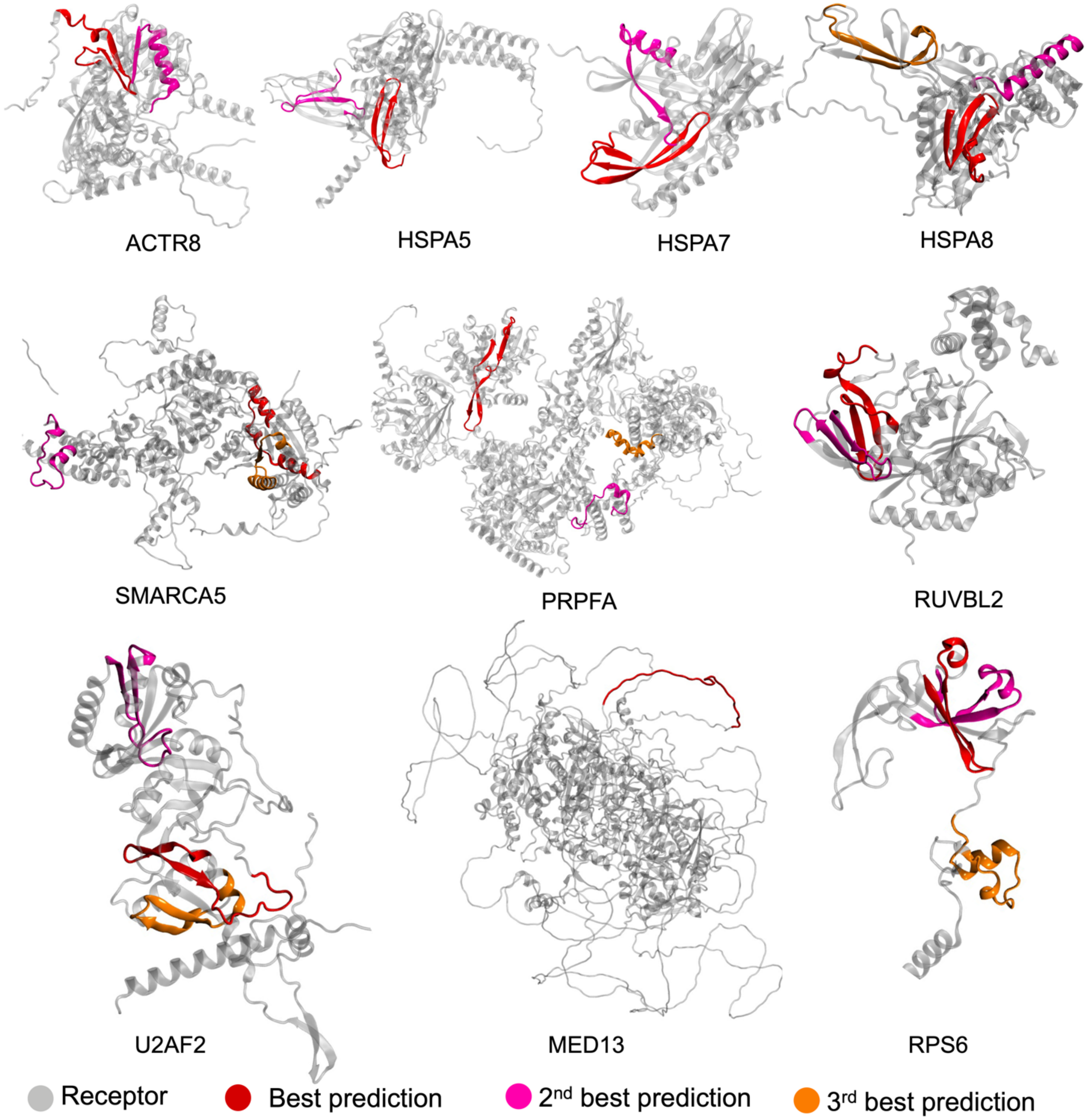


**Figure S23.** Full-length AF predicted structures of protein sequences derived from high-confidence pull-down hits. Top predicted epitope regions are shown red, magenta, and orange color. Rest of the proteins are shown in silver. AF regions that are not predicted have been observed to often correspond to intrinsically disordered regions in the protein.
